# Electronic cigarette vaping triggers lipid mediated vocal fold mucosal injury

**DOI:** 10.1101/2020.09.25.313486

**Authors:** Vlasta Lungova, Susan L. Thibeault

## Abstract

Electronic cigarettes (e-cigs) are nicotine delivery systems that have been touted as safer alternatives to smoking. A recently reported case of epiglottitis revealed a connection between vaping and swollen laryngeal and vocal fold (VF) structures that can lead to acute life-threatening airway obstruction. The clinical course and biopsy revealed direct epithelial injury and subsequent inflammatory reaction. Here we show that we were able to recapitulate this phenomenon in in vitro conditions. Exposure of engineered VF mucosae to 5% e-cig vapor extract for one week induced cellular damage in VF luminal epithelial cells, disrupting mucosal homeostasis and mucosal innate immune responses. Epithelial erosion was likely caused by the accumulation of solvents and lipid particles, most likely medium chain fatty acids, in the cytosol and intercellular spaces, which altered lipid metabolism and plasma membrane properties. In summary, vaping represents a threat to the VF mucosa health and airway protection.

## Introduction

Recent cases of acute electronic cigarette (e-cig), or vaping, associated lung injuries (EVALI) opened a debate about the safety and health-related consequences of vaping. Multiple case reports have described atypical pneumonia and deaths in e-cig users (1–5) prompting intense scientific research focusing on effects of vaping on cellular functions of distal airways, where gas exchange takes place (6–8). As e-cigs are heated in the mouth, inhaled vaporized e-liquids pass through the throat past the larynx and vocal folds (VF), down into the lungs. Local droplet deposition in upper airways can have, therefore, physiological consequences and pose potential threats to oropharyngeal and VF health. E-cigs are nicotine delivery systems have been touted as safer alternatives to conventional smoking and rapidly gained popularity, especially among young adults and high school age adolescents (9). E-cigs consist of prefilled or fillable cartridges with e-liquids that serve as reservoirs for vaping substances such as vehicle solvents, propylene glycol (PG) and vegetable glycerin (VG), mixed with different concentrations of nicotine (N) and flavors (F). When heated, these substances form an aerosol that is then inhaled. VG is thick with a natural sweet flavor, producing the clouds of vapors upon exhalation; PG is less viscous, producing greater throat stimulation and mimics the sensation of smoking. PG and VG give e-liquids their high viscosity. As a result, aerosols from these liquids are likely to adhere to exposed surfaces, such as the soft and hard tissues (10). Currently, an estimated 10 million US adults and over 3 million high school students are active e-cig users (9). Those that have no previous experience with conventional cigarette smoking are at higher risk, as they exhibit increased susceptibility to lung damage and viral and/or bacterial infections (11).

In this study, we evaluated possible consequences of vaping on VF mucosal structure and function. The larynx and VF are involved in voice production and are, as parts of the conducting airways, directly exposed to inhaled vaporized e-liquids. Particularly vulnerable are epithelial cells that serve as first line of defense against inorganic, organic, and microbial intruders and protect the lamina propria beneath (12). So far, vaping associated cases of respiratory diseases have sudden onset symptoms that develop rapidly (1–3,6). Acute diseases associated with the larynx include acute laryngitis and epiglottitis and are usually caused by viral or bacterial infections (13). They cause swelling of laryngeal structures and can rapidly lead to life-threatening airway obstruction (13). A recently reported case of epiglottitis in an adolescent female patient revealed a connection between vaping and swollen laryngeal structures without signs of viral, bacterial or fungal infections (14). She was hospitalized twice and presented with acute respiratory distress, severe dysphagia, hoarseness and increased throat clearing. She reported uses e-cigs over 1 to 2 months with different fruit- or candy-flavored cartridges. Stroboscopic examination showed moderate swelling of the epiglottis and supraglottal structures, pink laryngeal mucosa with marked inflammation. Her biopsies from the arytenoids and soft palate and a sample of fluid from the laryngeal/epiglottic region revealed reactive squamous epithelial changes with focal erosions, abundance of cellular debris and thick mucus. Her clinical course and biopsy findings were highly suspicious for direct chemical injury and/or subsequent inflammatory reaction (14). Despite the importance of understanding of the biological effect of vaping on laryngeal and VF mucosa, physiological consequences of e-liquid deposition and whether this phenomenon can be recapitulated in experimental *in vitro* conditions remains unknown.

The recent development of a three-dimensional (3D) model of human VF mucosa by our group has allowed us to mimic *in vivo* remodeling of the VF mucosa in tobacco-related diseases (15). Upon exposure to 5% cigarette smoke extract (CSE) for 1 week, we were able to induce keratotic changes and mucosal inflammation in engineered VF mucosae, composed of human induced pluripotent stem cell (hiPSC) derived VF epithelial cells and primary human VF fibroblasts dispersed in the collagen gel that mimicked the lamina propria. Here we demonstrate that exposure to 5% e-cig vapor extract (ECVE) for 1 week induces VF epithelial injury which compromises the integrity of apical cell layers, membrane-anchored luminal mucin production and clearance, and dysregulates VF mucosal immune responses. We further show that epithelial erosion is likely caused by the accumulation of vaporized lipophilic solvents and lipid particles, most probably medium-chain triglycerides, in the cytosol that alter the lipid metabolism and restructure plasma membrane properties.

Collectively, our experimental findings revealed that exposure of VF mucosa to vaporized e-liquids disrupts VF mucosal homeostasis and innate barrier functions, which represents a potential threat to the VF mucosa health, voicing and mainly airway protection, whereby raising concerns over the safety of e-cig use to other vital and essential portions of the upper airway.

## Results

### Histological alterations of VF mucosae exposed to 5% ECVE

To investigate the effect of ECVE on the human VF mucosa homeostasis, we utilized the recently developed hiPSC-derived model of human VF mucosa (15). HiPS cells were first differentiated into VF basal epithelial progenitors for 10 days and then reseeded on the top of collagen-fibroblast constructs and were let to differentiate for additional 22 days. At day 32, engineered VF mucosae were exposed to 5% ECVE for 1 week to mimic exposure of cells to vaping (Figure 1A) (for detailed protocol see the Material and Methods). VF mucosae treated with plain culture medium were used as negative controls. For experimental groups, we tested three different types of e-cig vapor extracts including vehicle controls with polypropylene glycol and vegetable glycerin (PG/VG) only, e-cigs with PG/VG and nicotine (PG/VG+N) and e-cigs with PG/VG+N and flavor (PG/VG+N+F), the most popular types of e-cigs. At Day 39, human engineered VF mucosae were collected for analysis.

**Figure 1:**
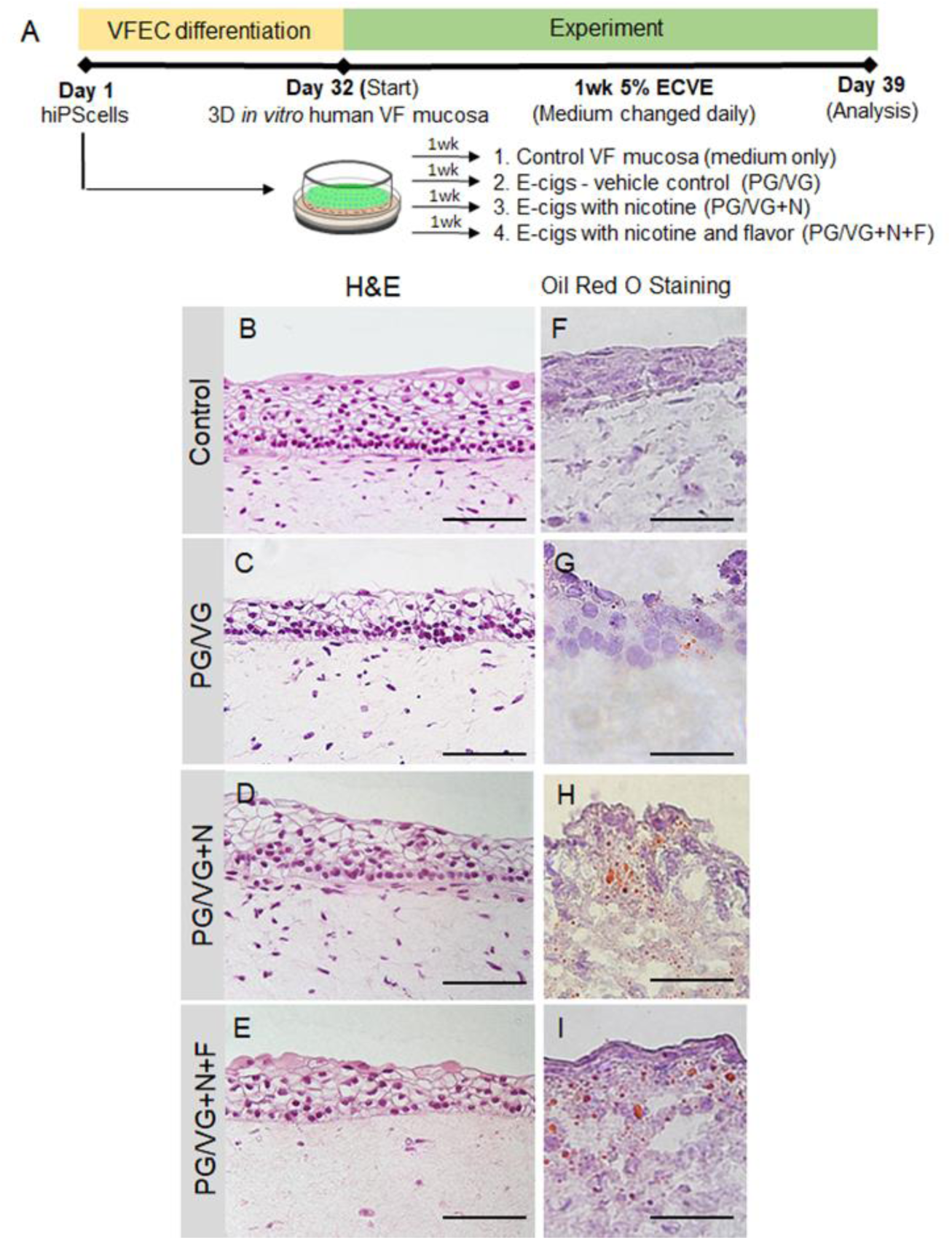
Experimental design and morphology of hiPSC-derived VF mucosae exposed to 5% ECVE. (A) Schematic illustration of the experimental design. hiPS cells were first differentiated into VF epithelial cells (VF EC) for 32 days and then exposed to 5% ECVE for 1 week. We tested three different conditions – e-cigs, vehicle controls, with PG/VG only, e-cigs with PG/VG and nicotine (PG/VG+N) and e-cigs with PG/VG with nicotine and flavor (PG/VG+N+F). Control VF mucosae treated with plain culture medium were used as negative controls. At Day 39, VF mucosae were collected and analyzed by immunohistochemistry and quantitative polymerase chain reaction. (B-E) Morphology of VF mucosae in control group (B) and 5% ECVE exposed groups (C-E) showing stratified squamous VF epithelium. (F-I) Oil Red O stain on frozen unfixed sections of VF mucosae in the control group (F) and 5% ECVE exposed groups (G-I). All 5% ECVE treated samples contained lipid droplets that adhered to cell surfaces. Scale bar = 100 μm (B-E) and 50μm (F-I). Abbreviations: e-cigs, electronic cigarettes; ECVE, electronic cigarette vapor extract; hiPS, human induced pluripotent stem cells; VF, vocal folds; VFEC, vocal fold epithelial cells.

First, we investigated alternations in morphology of VF mucosae in control and experimental groups. Histological assessment of the VF mucosae showed a typical stratified squamous architecture for all conditions (Figure 1B-E). Epithelial cells were arranged in a basal cell layer and suprabasal cell layers that flattened apically (Fig. 1B-E). Gross histological examination revealed that ECVE exposed VF mucosae appeared thinner than control (Figure 1B-E). Next, we performed Oil Red O staining on frozen unfixed sections and found red oil droplets on the surface of the VF epithelium in all ECVE exposed VF mucosae (Figure 1F-I), suggesting that e-liquids contained lipid components. When heated, e-liquids produced aerosol mixing with the culture medium, and depositing lipid droplets that adhered to cell surfaces. In order to examine whether the exposure of cells to ECVE could alter the structure and function of the VF epithelial barrier we evaluated expression of stratified epithelial markers and tested functionality of VF epithelial cells by assessing mucin and inflammatory cytokine/chemokine expression.

### ECVE affects compactness of apical epithelial cell layers

We have previously demonstrated that exposure of engineered VF mucosae to 5% cigarette smoke extract (CSE) leads to VF mucosa remodeling affecting predominantly the basal epithelial cell layer with downregulation of cytokeratin (K) 14 which pathologically accumulates in the luminal cells along with K13 (15). Therefore, we sought to determine whether 1-week exposure to 5% ECVE could also affect cytokeratin production and structure of the basal cellular compartment. We first evaluated K14 and Laminin alfa 5 (LAMA5) expression and co-stained with p63. We found that K14 did not change in 5% ECVE treated groups versus controls (Figure 2A-D). Similarly, LAMA5, a marker of the basement membrane, was detected in both control and 5% ECVE treated groups (Figure 2E-H) along with p63+, a marker of basal cells (Figure 2 A-H). However, there was a reduced expression of suprabasal K13 in ECVE exposed groups, as compared to controls suggesting that apical epithelial surfaces are compromised by ECVE (Figure 2I-L). To assess the compactness of the epithelial barrier, we stained for E-Cadherin (E-Cad), a marker of cell adherent junctions. We found that in control samples, E-Cad was strongly expressed in all epithelial cell layers (Figure 2M) while in 5% ECVE exposed groups E-Cad signal was absent in some cells in the apical cell layers (Figure 2N-P). Histological data were supported by qPCR (Figure 2Q-T). As expected, transcript levels of K14 did not change in the PG/VG+N+F group or were upregulated in the PG/VG and PG/VG+N groups as compared to control (Figure 2Q). p63 remained the same in PG/VG and PG/VG+N groups and significantly decreased in PG/VG+N+F (Figure 2R). On the other hand, unlike in control, expression levels of K13 significantly dropped down in ECVE exposed groups (Figure 2S). A similar pattern was observed in E-Cadherin (encoded by CDH1 gene), which significantly decreased in all experimental groups compared to controls (Figure 2T). These data indicate that exposure of the VF epithelium to 5% ECVE likely impairs the structure and integrity of the luminal epithelial cell layers that come into the direct contact with aerosol and toxic substances found in ECVE. Moreover, the decreased expression of p63 in the PG/VG+N+F group suggests that the damage of the luminal layers can also ultimately cause changes in basal cells.

**Figure 2:**
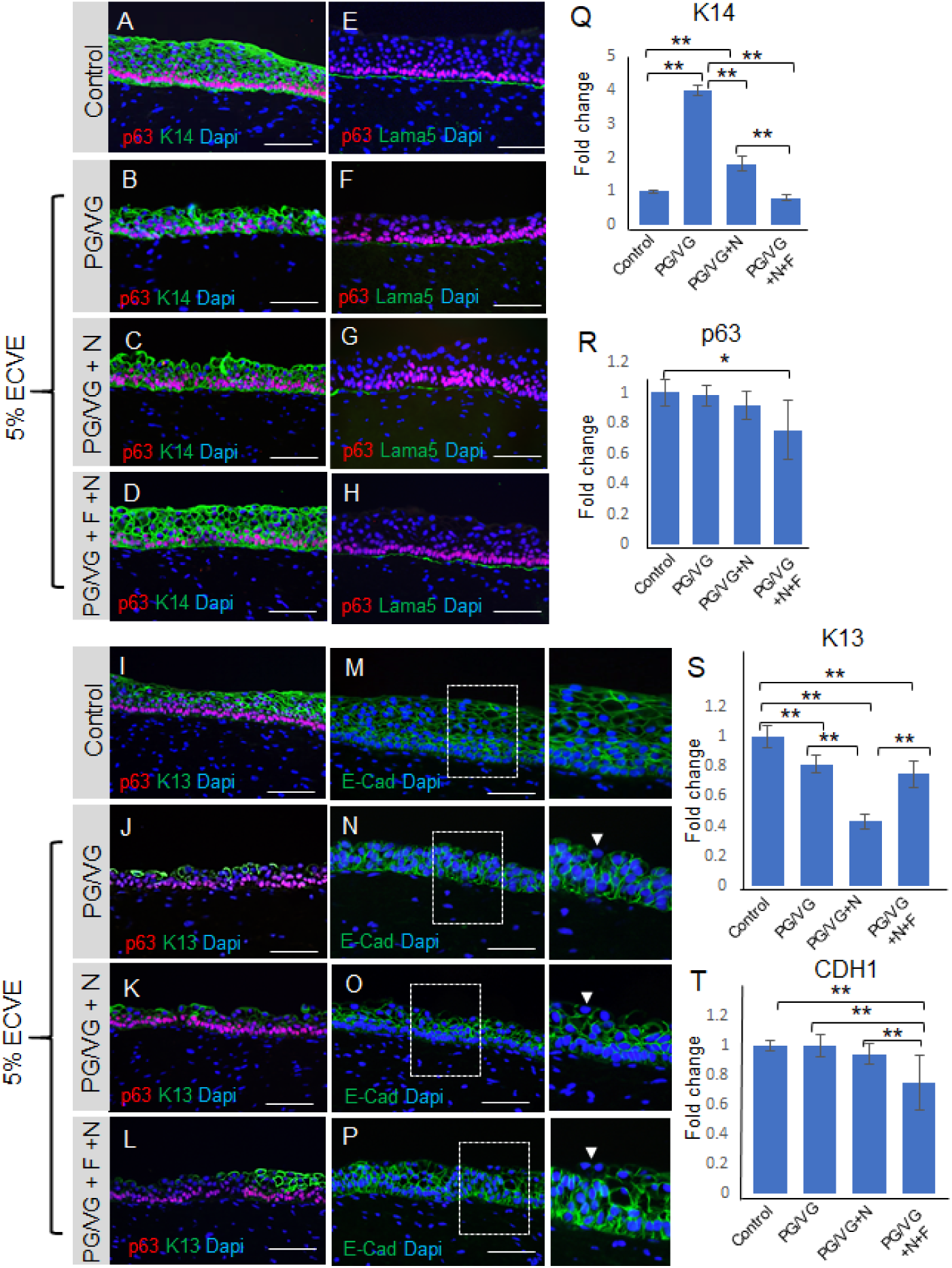
Distribution and expression levels of VF structural epithelial genes. (A-D) Anti-Cytokeratin K14 (green) and anti-p63 staining (red) in control (A) and 5% ECVE exposed VF mucosae (B-D). (E-H) Anti-Laminin alfa 5 (green) co-stained with anti-p63 (red) in control (E) and 5% ECVE treated VF mucosae (F-H). (I-L) Anti-cytokeratin K13 (in green) co-stained with anti-p63 (in red) in control (I) and 5% ECVE exposed VF mucosae (J-L). (M-P) Anti-E-cadherin staining (in green) in control (M) and 5% ECVE exposed VF mucosae (N-P). Bracketed regions in panels M, N, O and P are magnified in the boxes on the right. White arrow heads point to apical cells with decreased expression of E-Cadherin. Scale bars = 100μm. (Q-T) Transcript levels of cytokeratin K14 (Q), p63 (R), K13 (S) and CDH1 (T) in control and 5% ECVE exposed VF mucosae. Error bars represent ± standard error of the mean obtained from two biological and three technical replicates. One-Way ANOVA of variance for independent or correlated samples statistical analysis along with Tukey HSD test were used to confirm statistical significance in gene expression {p-Value ≤ 0.05 (*) and p-Value ≤ 0.01 (**)].

### ECVE alters membrane-associated mucin and cytokine expression

We further investigated whether exposure to 5% ECVE can alter the function of the VF epithelial protective barrier. We have previously shown that the VF epithelium is an essential mechanism of VF defense (12) which is achieved by the compact physical epithelial barrier, mucus production and secretion of cytokines/chemokines that are parts of the innate immunity. Above, we have shown that 5% ECVE exposure affects cell adherent junctions and K13 expression in the luminal cell layers. In this section, we will primarily focus on the mucin and cytokine/chemokine expression. We evaluated expression of Mucin1 (MUC1) and 4 (MUC4) which are typical membrane-associated mucins present in the human VF (12, 15). Secretory proteins, such as MUC5B or MUC5C, are not produced by stratified VF epithelial cells that cover the membranous portion of the true VF (12). Our histological data confirmed that in control VF mucosae MUC1 was detected in the apical cell layer and formed a thin protective coat (Figure 3A) as previously shown in the human native VF mucosae (15). In 5% ECVE treated groups PG/VG and PG/VG+N, MUC1 expression appears to be upregulated and moved into deeper epithelial cell layers (Figure 3B, C). Notably, in the PG/VG+N+F group, the MUC1 layer remained thin, but we observed mucus clots on the epithelial surface that wrapped cell debris (Figure 3D). As for MUC4, we did not observe any significant changes in the expression pattern between control and ECVE exposed groups (Figure 3E-H). We further confirmed expression of mucins by qPCR. In PG/VG and PG/VG+N, MUC1 transcript levels were upregulated, while in PG/VG+N+F group the expression levels of MUC1 were similar to controls (Figure 3I). On the other hand, transcript levels for MUC4 decreased particularly in the PG/VG+N and PG/VG+N+F exposed VF mucosae (Figure 3I). These findings show that exposure to ECVE leads to the increased expression of luminal MUC1, but not MUC4, and formation of mucus clots that can accumulate in the laryngeal/airway lumen and impair mucus clearance.

**Figure 3:**
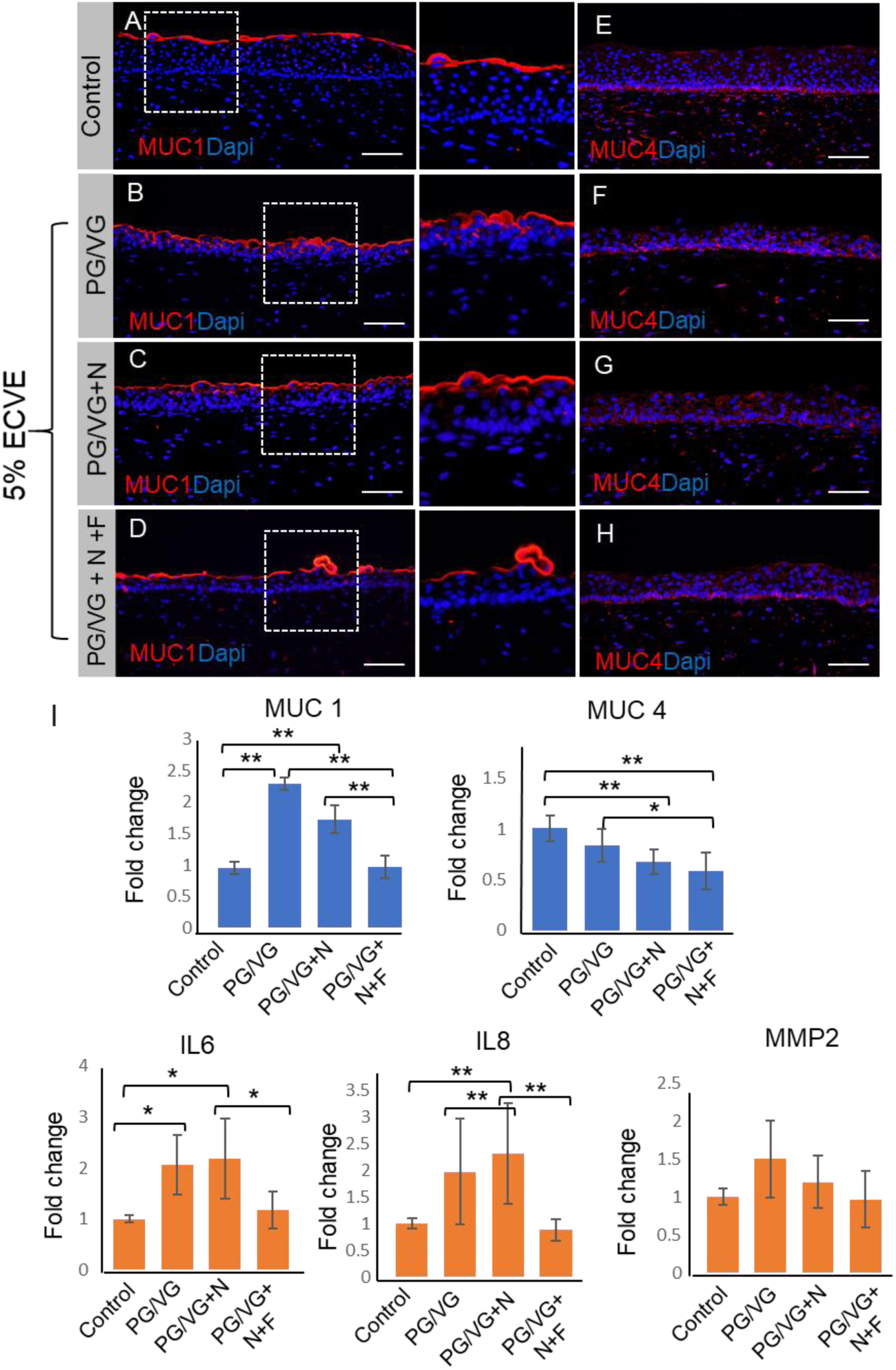
Distribution and expression levels of functional VF epithelial genes. (A-D) Red immunofluorescent staining of Mucin 1 (MUC1) in control (A) and 5% ECVE exposed VF mucosae (B-D). Bracketed regions in panels A-D are magnified on the right respectively. (E-H) Red immunofluorescent staining of Mucin 4 (MUC4) in control (E) and 5% ECVE treated VF mucosae (F-H). Scale bar = 100μm. (I) Transcript levels of MUC 1 and MUC 4 and selected genes involved in the VF immune response - nterleukin IL6, IL8 and matrix metalloproteinase 2, MMP2. Error bars represent ± standard error of the mean obtained from two biological and three technical replicates. One-Way ANOVA of variance for independent or correlated camples statistical analysis along with Tukey HSD test were used to confirm statistical significance in gene expression {p-Value ≤ 0.05 (*) and p-Value ≤ 0.01 (**)].

Next, we assessed whether structural and functional changes in the VF epithelium were capable of inducing expression of cytokines and activate a VF mucosal immune response (Figure 3I). Although we found upregulation of interleukins (IL) IL6 and IL8 in ECVE exposed groups, particularly in PG/VG and PG/VG+N groups in comparison to controls, expression levels varied and were lower than previously reported in 5% CSE exposed VF mucosae (15). Moreover, MMP2 expression did not change in control versus 5% ECVE exposed groups (Figure. 3I). We sought to further elucidate, whether this moderate inflammatory response involves selected cytokines only or whether suppression of the mucosal immune response is more general affecting cell-cell signaling, chemotaxis and rapid response of cells to infectious agents.

### ECVE stimulates production of chemokines implicated in the recruitment of eosinophils

To better understand the immunomodulatory consequences of ECVE exposure, we next evaluated VF mucosa cytokine and chemokine profiles from control versus ECVE exposed VF mucosae. We performed SYBR green-based quantitative real-time RT^2^ PCR profiling array looking at genes involved in Human Cytokine and Chemokine Expression. We used 384-well profiler plates that were designed to screen all four samples on one plate (a 4 x 96-well format). Samples were run in a triplicate (n=3; 12 samples total). For each sample, the 384-well plate contained primers for 84 genes involved in human cytokine and chemokine expression, 5 housekeeping genes and 3 negative control wells (Supplemental Table S1). CT values, fold-regulation and p-Values for all tested genes are shown in Supplemental Tables S2 and >S3. Gene expression levels were normalized to reference (housekeeping) genes and control samples. Positive fold-regulation values indicate upregulated genes and are equivalent to the fold change. While negative values indicate downregulated genes and are negative inverse of the fold change (Supplemental Table S3). Genes with a fold change ≥ 2 and p-Value ≤ 0.05 were considered as significantly differentially expressed genes. Our results revealed that exposure of the VF mucosae to 5% ECVE dysregulated inflammatory responses in epithelial cells and VF fibroblasts (Figure 4). In control versus PG/VG groups, we found significant modulation in 7 genes (Figure 4A), with 2 upregulated genes BMP4 (2.89-fold) and BMP7 (2.49-fold), and 5 downregulated genes *SPP1* (-3.18-fold), *IL23A* (-8.35-fold)*, IL21* (-7.39-fold)*, CXCL12* (-3.83-fold) *and CCL7* (-3.88-fold). In the PG/VG+N group we found 5 genes significantly different from controls (Figure 4B) with 1 gene upregulated *CCL11* (2.93-fold) and 4 suppressed genes including *THPO* (-2.54-fold)*, IL4* (-3.77-fold)*, IL23A* (-3.98-fold) and *CSF3* (-2.94-fold). When we compared the PG/VG+N+F group with the controls, 12 genes showed significant differential expression (Figure 4C); *IL6* (3.01-fold) and *CCL11* (2.97-fold) were upregulated and *SPP1* (-7.17-fold)*, MSTN* (-2.59-fold)*, IL4* (-4.10-fold)*, IL23A* (- 9.10-fold)*, IL21 (-11.12-fold), CX3CL1* (-2.89-fold)*, CSF3* (-2.76-fold)*, CCL5* (-4.87-fold)*, BMP4* (-2.22-fold) and *ADIPOQ* (-4.34-fold) were downregulated. These data suggest that the combination of PG/VG+N+F dysregulates VF mucosal cytokine production more than with PG/VG and PG/VG+N. Suppression of chemokines, such as *CCL5, CCL7, CX3CL1, IL23A, IL21 and CSF3,* may cause the delay in mucosal VF responses to pathogens (7), as these chemokines are implicated in recruitment of monocytes/macrophages and neutrophils to the site of inflammation (16, 17). Suppression of cytokines with anti-inflammatory cytoprotective function, such as *IL4* (18) may slow down the VF mucosal repair and regeneration. On the other hand, consistent elevated transcript levels of *CCL11* indicate that ECVE can activate migration of eosinophils that are otherwise involved in allergic reactions (19). The differences in the secretion of cytokines/chemokines suggest that ECVE likely impairs innate epithelial barrier function along with VF fibroblast immune responses, which can lead to the development of allergic inflammation and/or decreased protection and recovery time of the VF epithelial barrier.

**Figure 4:**
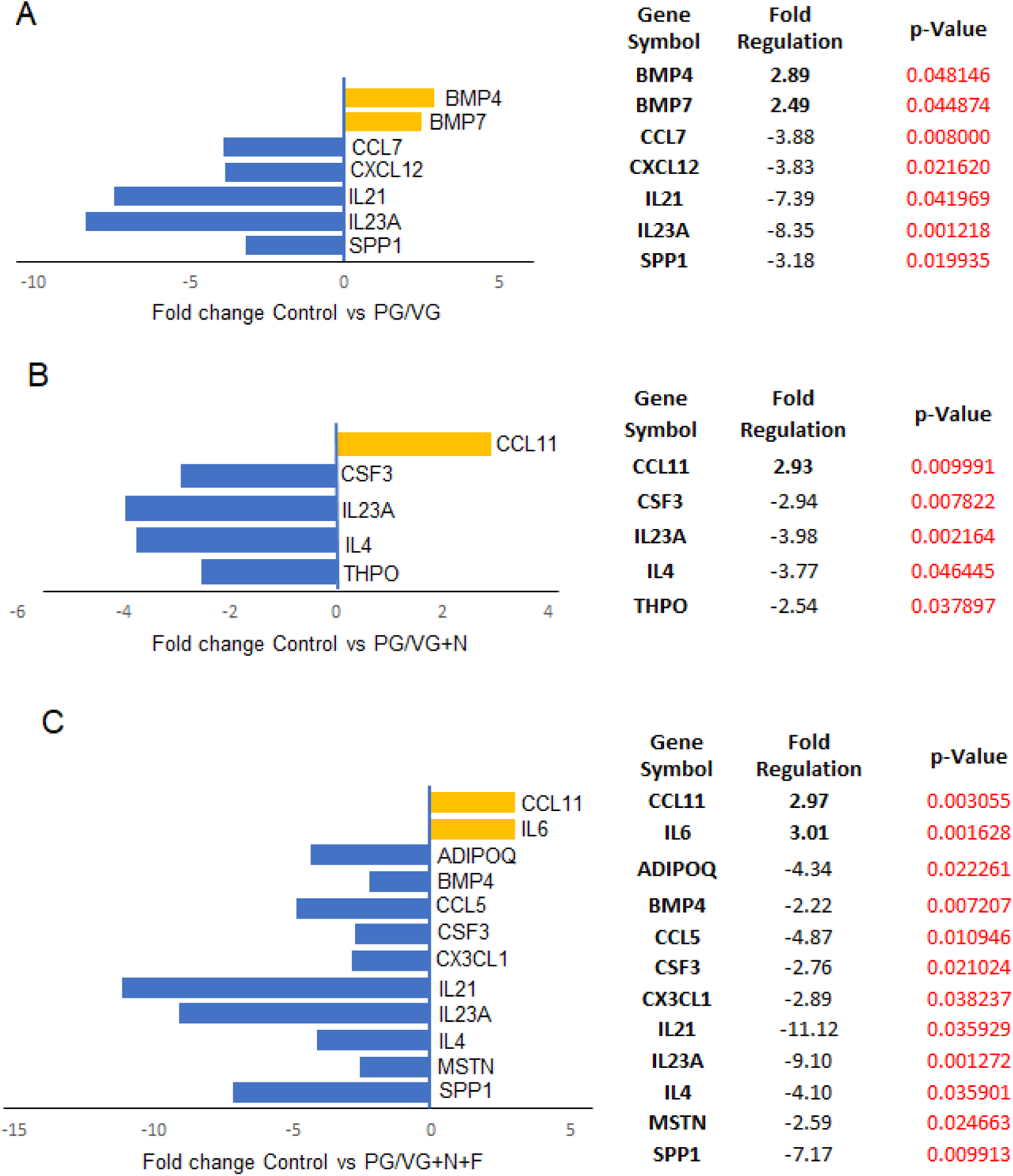
RT^2^ PCR profiling analysis focusing on human cytokine and chemokine expression. (A-C) Significantly differentially expressed genes involved in mucosal inflammation in control versus PG/VG group (A), control versus PG/VG and nicotine (B) and control versus PG/VG and nicotine and flavor (C). Upregulated genes with positive fold-regulation values are highlighted in yellow color. Downregulated genes with negative fold-regulation values are highlighted in blue. The fold-change threshold was set to 2. P-values were calculated based on a Student’s t-test of the replicate 2∧(-Delta Delta CT) values for each gene in the control group and treatment groups, and p-values less than 0.05 were considered as significant. P-value calculation used was based on parametric, unpaired, two-sample equal variance, two-tailed distribution. Samples were run in a triplicate (n=3, 12 samples total).

### ECVE exposure causes imbalance in lipid cytosolic content and alters plasma membrane properties

We further investigated whether altered VF epithelial structure and function can be associated with oil droplet deposition and examined the ultrastructure of VF epithelial cells with transmission electron microscope (TEM). In ECVE exposed VF mucosae, but not in controls, we found numerous of tiny dark spots (more than likely droplets) along with large white lipid aggregates, inside the cytoplasm and intercellular spaces (Figure 5A-H). The presence of inclusions between plasma membranes dilated the space between neighboring cells and impaired cell junctions (Figure 5C). Moreover, in the PG/VG +N and PG/VG+N+F groups, outermost epithelial cells detached from the underlying cell layers (Figure 5E, G), some exfoliated cells became loaded with lipid particles and dark pigments (Figure 5G, H) and resembled lipid laden macrophages (6, 7). The origin of the cytoplasmic lipid aggregates and/or dark pigments is not clear. Since VF epithelial cells are not active producers of lipid components, such as pulmonary surfactants (7), we speculate that these are from exogenous sources. Despite the fact that the VF epithelial cells are not active lipid producers they possess the enzymatic machinery involved in lipid/fatty acid metabolism to maintain phospholipid by-layers and vital cellular functions. To investigate whether the cytoplasmic accumulation of lipid droplets can affect VF mucosal cell metabolic activity related to fatty acid/glycerol breakdown and/or recycling we performed SYBR green-based quantitative real-time RT^2^ PCR profiling array looking at genes involved in Human Fatty Acid Metabolism. As described above, a 4 x 96-well format (384-well plates) was used to screen all four samples on one plate and data were run on three separate plates (n=3, 12 samples total). For each sample, the 384-well plate contained primers for 84 genes involved in human fatty acid metabolism, 5 housekeeping genes and 3 negative control wells (Supplemental Table S4). CT values, fold-regulation values and p-Values for all tested genes are included in the Supplemental Tables S5 and S6. As mentioned above, positive fold-regulation values indicate upregulated genes and are equivalent to the fold change, while negative fold regulation values indicate downregulated genes and are negative inverse of the fold-change (Supplemental Table S6). Genes with a fold change ≥ 2 and p-Value ≤ 0.05 were considered as significantly differentially expressed genes.

**Figure 5:**
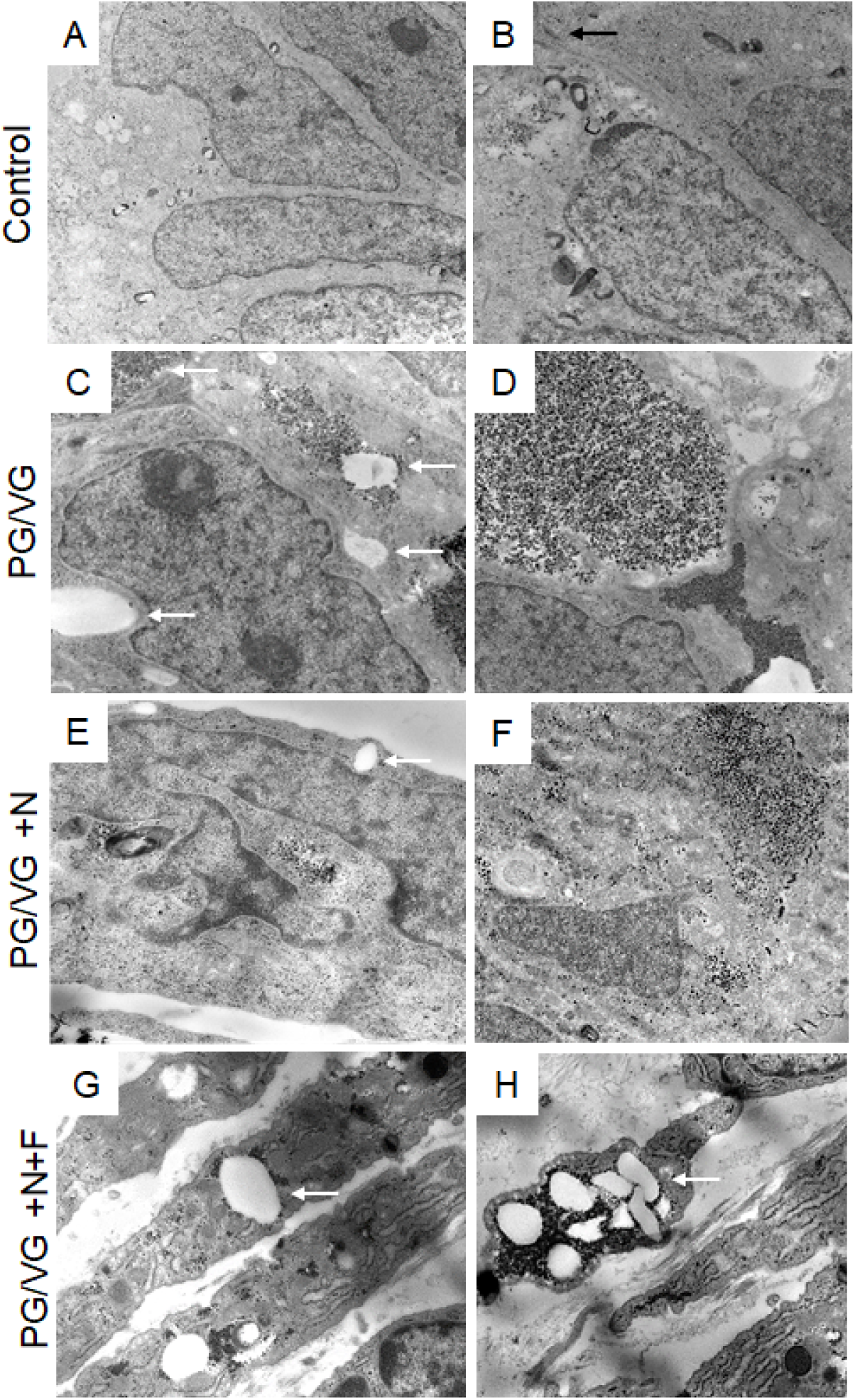
Transmission electron microscopy of the apical region of the VF epithelium. (A, B) Ultrastructure of VF epithelial cells of the control VF mucosa. (C, D) Ultrastructure of VF epithelial cells exposed to PG/VG only. Cytosol of the epithelial cells contains dark droplets and/or pigments. White solid arrows denote white aggregates that accumulate in the cytosol and intercellular space. (E, F) Ultrastructure of the VF epithelial cells exposed to PG/VG and nicotine. White solid arrows point to white lipid aggregates in the cell cytoplasm. Apical cells tend to detach from the underlying epithelial cell layers (G, H) Ultrastructure of the VF epithelial cells exposed to PG/VG and nicotine and flavor. White solid arrows point to white lipid aggregates and dark droplets in the cell cytoplasm of detached cells. Some exfoliated cells became loaded with lipid particles and dark pigments and resembled lipid laden macrophages.

**Figure 6:**
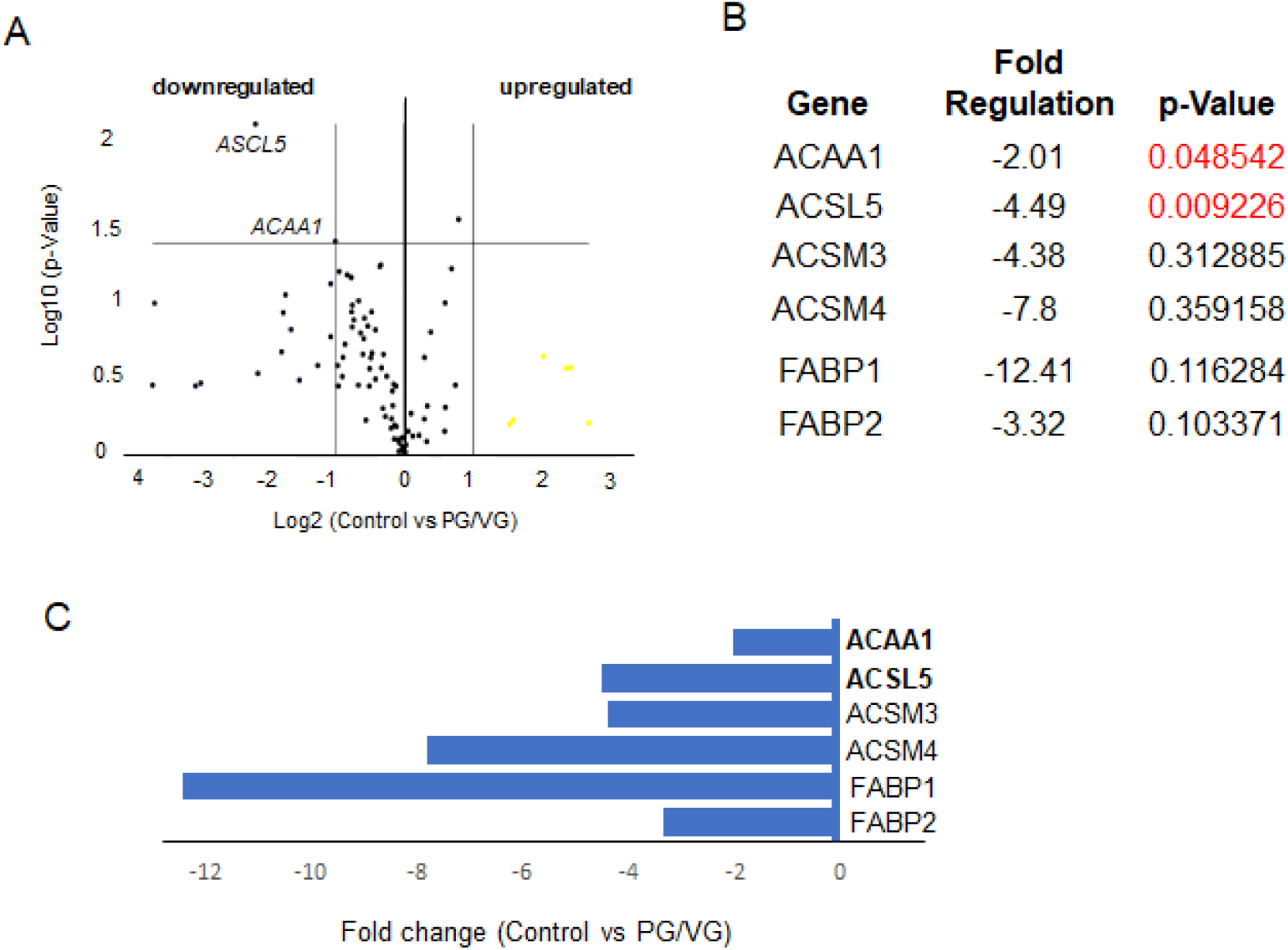
RT^2^ PCR profiling analysis focusing on human fatty acid metabolism in control and PG/VG exposed VF mucosae. (A) Volcano Plot identifies significant gene expression changes by plotting the log2 of the fold changes in gene expression on the x-axis versus their statistical significance on the y-axis. The center vertical line indicates unchanged gene expression, while the two outer vertical lines indicate the selected fold regulation threshold: fold-change ≥ 2. The horizontal line indicates the selected p-value threshold ≤.0.05. (B) A table showing the list of differentially expressed genes in control versus PG/VG test group with fold-regulation values and p-values. P-Value highlighted in red color indicate significantly differentially expressed genes with fold change greater than 2 and a p-Value less than 0.05. (C) Graph showing differentially expressed genes in control versus PG/VG test group with fold-regulation plotted on the x-axis. Downregulated genes are highlighted in blue.

Our results show that the PG/VG group was at least affected by 5% ECVE as only 2 genes were significantly different from controls. ACAA1 (-2.01-fold) and ACSL5 (-4.49-fold) (Figure 6A-C) were significantly downregulated (Figure 6B). In other experimental groups, PG/VG+N and PG/VG+N+F, the effect of ECVE was more obvious (Figures 7 and 8). We found 9 and 5 significantly differentially expressed genes in PG/VG+N and PG/VG+N+F, respectively. Upregulated genes included ACADM with a fold change of 32.85 in PG/VG+N and 9.59 in PG/VG+N+F (Figures 7 and 8). This gene encodes the enzyme, Acyl-Coenzyme A dehydrogenase, that works on medium-chain fatty acids and is involved in medium-chain fatty acid degradation in mitochondria (20). On the other hand, downregulated genes, ACSBG2 (-5.38-fold), ACSL5 (-3.12-fold, ACSM5 (-3.62-fold), CPT1B (-2.98-fold) in PG/VG+N group and ACSBG2 (-3.58-fold), ACSM5 (-3.07-fold) in PG/VG+N+F group (Figures 7 and 8) encode enzymes needed for the synthetic reaction of fatty acids with acyl-Coenzyme A and adenosine triphosphate in the cytosol which is required for fatty acid activation and their translocation into mitochondria (21, 22). Notably, significantly downregulated ACSM5 in PG/VG+N and PG/VG+N+F and other downregulated members of the family, such as ACSM4 and ACSM3 (Figure 7 and 8) also work on medium-chain triglycerides/fatty acids (22). Moreover, we also found a significant downregulation of transport *FABP,* fatty acid binding proteins, particularly *FABP7*, with -3.67-fold in PG/VG+N and -2.18-fold in PG/VG+N+F; along with *SLC27A5* with -3.60-fold in PG/VG+N. These genes facilitate the transport of fatty acids/triglycerides across the plasma membrane and within the cytoplasm and thus regulate lipid cytoplasmic content (23).

**Figure 7:**
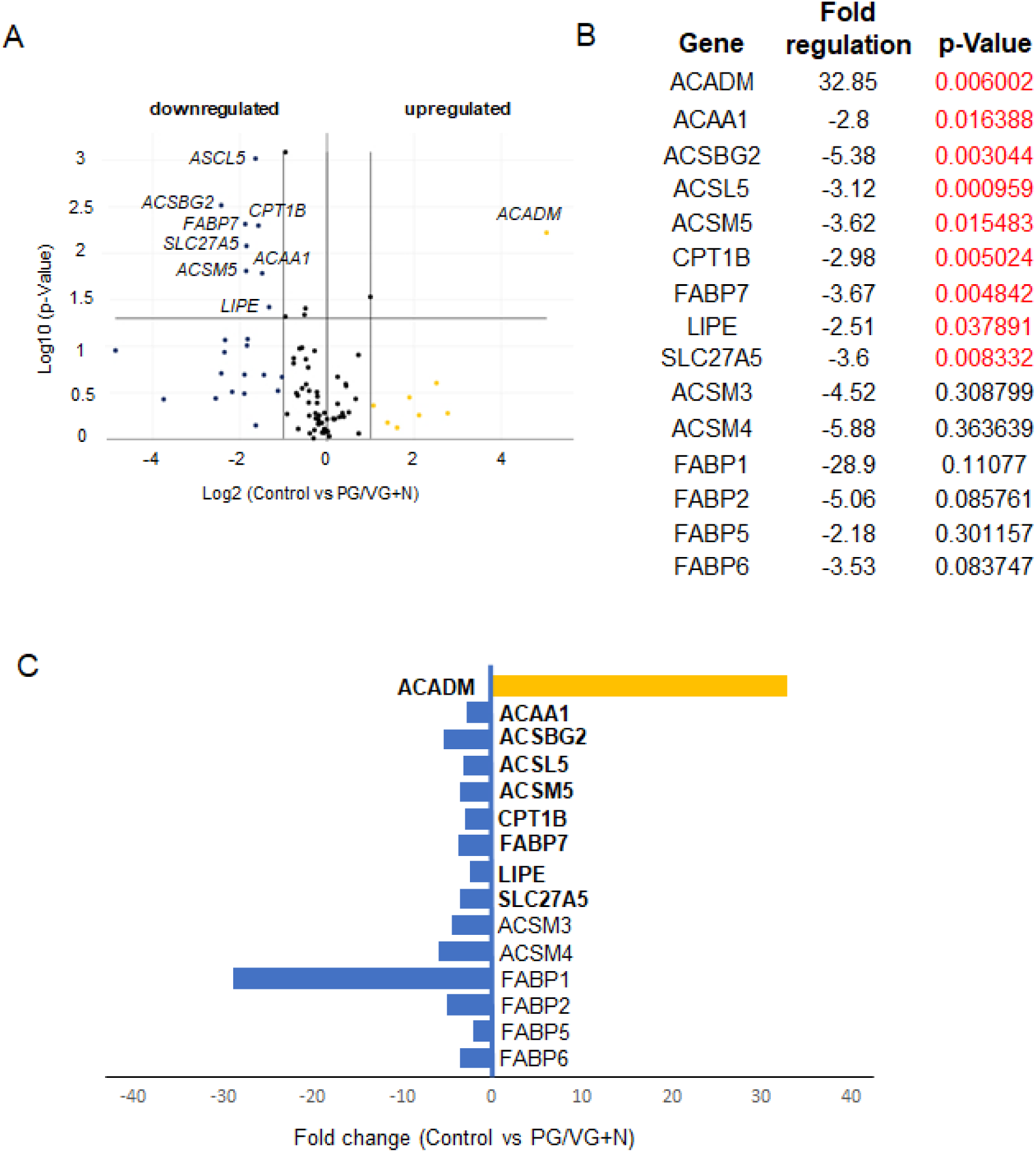
RT^2^ PCR profiling analysis focusing on human fatty acid metabolism in control and PG/VG and nicotine exposed VF mucosae. (A) Volcano Plot identifies significant gene expression changes by plotting the log2 of the fold changes in gene expression on the x-axis versus their statistical significance on the y-axis. The center vertical line indicates unchanged gene expression, while the two outer vertical lines indicate the selected fold regulation threshold: fold-change ≥ 2. The horizontal line indicates the selected p-value threshold ≤.0.05. (B) Table showing the list of differentially expressed genes in control versus PG/VG and nicotine test group with fold-regulation values and p-values. P-Values highlithed in red color indicate significantly differentially expressed genes with fold change greater than 2 and a p-Value less than 0.05. (C) Graph showing differentially expressed genes in control and PG/VG and nicotine test group with fold-regulation plotted on the x-axis. Upregulated genes are highlighted in yellow, downregulated genes are highlighted in blue.

**Figure 8:**
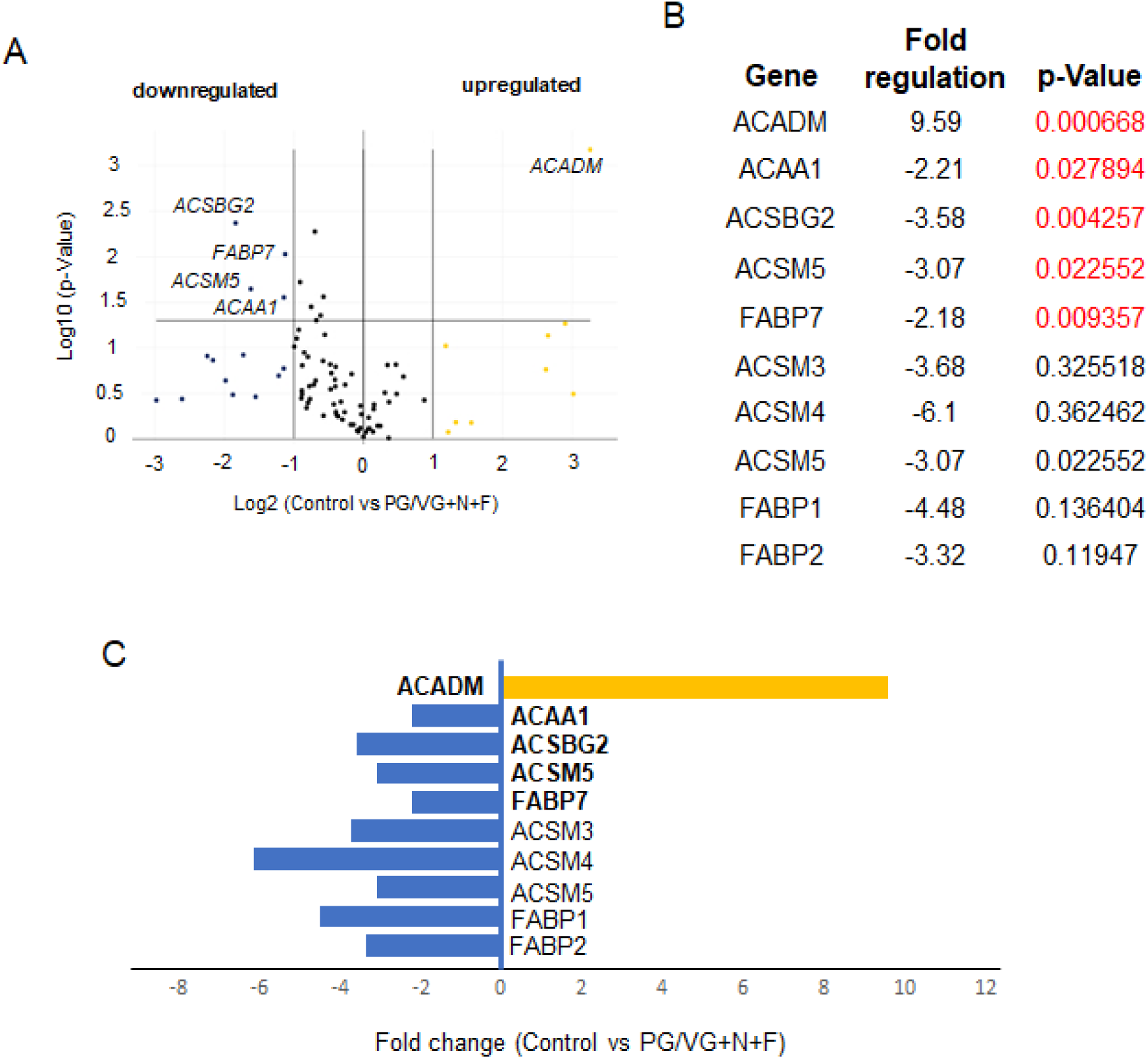
RT^2^ PCR profiling analysis focusing on human fatty acid metabolism in control and PG/VG and nicotine and flavor exposed VF mucosae. (A) Volcano Plot identifies significant gene expression changes by plotting the log2 of the fold changes in gene expression on the x-axis versus their statistical significance on the y-axis. The center vertical line indicates unchanged gene expression, while the two outer vertical lines indicate the selected fold regulation threshold: fold-change ≥ 2. The horizontal line indicates the selected p-value threshold ≤.0.05. (B) A table showing the list of significantly expressed genes in control and PG/VG and nicotine and flavor test group with fold-regulation values and p-values. P-Values highlighted in red color indicate significantly differentially expressed genes with fold change greater than 2 and a p-Value less than 0.05. (C) Graph showing differentially expressed genes in control and PG/VG and nicotine and flavor test group with fold-regulation plotted on the x-axis. Upregulated genes are highlighted in yellow, downregulated genes are highlighted in blue.

Overall, these findings show that the inefficient clearance of lipid and solvent substances in the cytosol dysregulates the lipid metabolism and plasma membrane properties. Besides in cytoplasm, lipid particles also deposit in the intercellular spaces impairing the integrity of the epithelial barrier with the secondary effect on communication between neighboring cells and cell-immune system signal transduction.

## Discussion

Our recently developed model of human VF mucosa holds great promise for studying toxicity and mechanisms of vaping-related cellular injuries in human VF stratified epithelium. Here we show that exposure of cells to 5% ECVE for one week was sufficient to induce cellular damage in VF apical epithelial cells, which disrupted the VF mucosal homeostasis and innate barrier function. Our results correlate with clinical observations (14), as well as results published for other types of epithelia, most notably, airway and nasal epithelia (7,25–27), suggesting that responses of cells to ECVE could be both, universal and tissue specific. Commonality with other studies includes compromised mucociliary clearance, immune responsiveness and aberrant lipid homeostasis. Among e-cig users, a significant increase in membrane-anchored mucins, and an increase in the ratio of secretory mucins MUC5AC to MUC5B has been reported compared to non-smoking participants (25). Nasal scrapes from e-cig users have shown significantly decreased expression of early growth response markers essential for the host-defense mechanisms as compared to cigarette smokers (26). Histopathological examination of lung biopsies and bronchoalveolar lavage (BAL) obtained from patients with EVALI revealed pigmented lipid laden macrophages and cell debris in the BAL fluid samples (5,6,27). In animal studies, mice receiving e-cig vapor exposure for 4 months failed to develop pulmonary inflammation and emphysema and exhibited delayed but enhanced response to the infectious agents (7). Moreover, suppression of the immune system was accompanied with altered lipid homeostasis both in resident alveolar macrophages and epithelial cells that secrete pulmonary surfactants (7). Specific to laryngeal and VF tissue, in a rat model, exposure to e-cig vapors for 4 weeks caused mild squamous metaplasia and hyperplasia and moderate subepithelial inflammation (24). However, differences between control and experimental animals were not statistically significant. Moreover, histological VF transversal sections of e-cig exposed VF exhibited cell debris that accumulated in the laryngeal lumen. These findings correspond with our observations.

In this study, comprehensive genomic and structural analyses of human VF mucosal cells show that exposure of cells to 5% ECVE disrupts the apico-basal polarity of the VF stratified squamous epithelium. Under healthy steady-state conditions, the basal cellular compartment firmly anchors the VF epithelium to the LP and provides the reserve of cells necessary for self-renewal (15, 28), while apical differentiated cell layers face the external environment and perform specific functions, most notably barrier formation (12), mucus secretion (29), sensory transduction and immunological surveillance (30, 31). Both domains form the compact VF epithelium held together by cell junctions that provide structural support and seal intercellular spaces (12). Induced epithelial injury, that removes apical cell layers, compromises the function of the entire epithelial protective barrier and exposes basal proliferating cells to further damage and pathogen infiltration (Figure 9). Detached cells wrapped in mucus accumulate in the airway lumen, which may contribute to increased throat clearing and coughing in e-cig users (11, 14). Our findings also show moderate inflammation in VF mucosae treated with 5% ECVE, which is in accordance with published data (7,26,32). Nevertheless, we evaluated cytokine/chemokine expression in VF mucosal cells without the presence of immune cells and macrophages, which could influence their transcript levels. Further validation of our 3D *in vitro* system, and cultivation of VF mucosal cells with a population of immune cells, will be extremely useful to provide a profound analysis of the effect of ECVE on VF mucosal inflammation.

**Figure 9:**
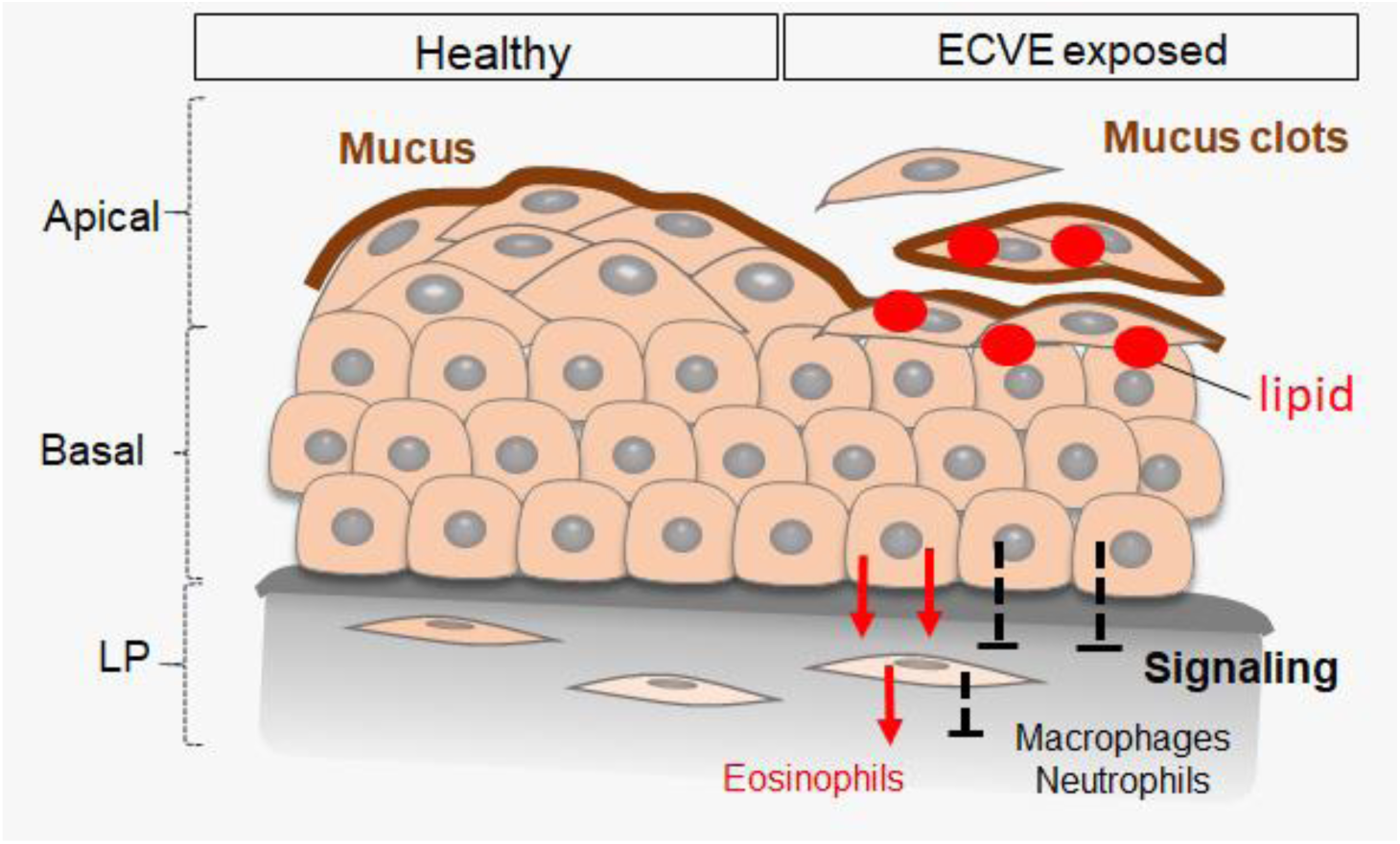
Schematic illustration showing the effect of 5% ECVE on the vocal fold stratified epithelium. Vocal fold epithelium on the left represents healthy stratified squamous epithelium with the mucus protective layer that covers the luminal epithelial surface. Vocal fold epithelium on the right side represents VF epithelium exposed to 5% PG/VG and nicotine and flavor. Due to the lipid droplet deposition superficial cell layers detach form the underlying cell layers and are wrapped in mucus. Removal of apical cell layers disrupts the signaling between the VF epithelial cells and fibroblast with immune cells. We detected upregulation of genes involved in recruitment of eosinophils, but not macrophages or neutrophils, which can lead to the development of allergic inflammation and/or decreased protection and recovery time of the vocal fold epithelial barrier. Abbreviations: Lr, larynx; OC, oral cavity; Ph, pharynx; Tr, trachea, VF, vocal folds.

The proposed mechanism underlying structural and functional changes of VF epithelium is likely associated with defective lipid metabolism and excess lipid/solvent particles that accumulate in the cytosol and intercellular spaces as confirmed by Red O stain and TEM. Lipid particles can be derived from nicotine and flavorings added to e-liquid to intensify the taste and/or enhance vaping experience (8). We found significant upregulation of ACADM in the VF mucosae exposed to commercially available e-liquids with nicotine and nicotine and flavor supporting the fact that lipid aggregates found in the cell cytoplasm likely contain medium-chain fatty acids/triglycerides that pass through the cell membrane by passive diffusion or via carriers linked to FABP proteins (23). Increased cytosolic deposition of these fatty acids is likely caused by downregulation of Acyl-CoA synthetase medium-chain family members - ACSM5, ACSM4 and ACSM3 that fail to activate fatty acids and prevent their translocation into mitochondria for final degradation (Figure 10).

**Figure 10:**
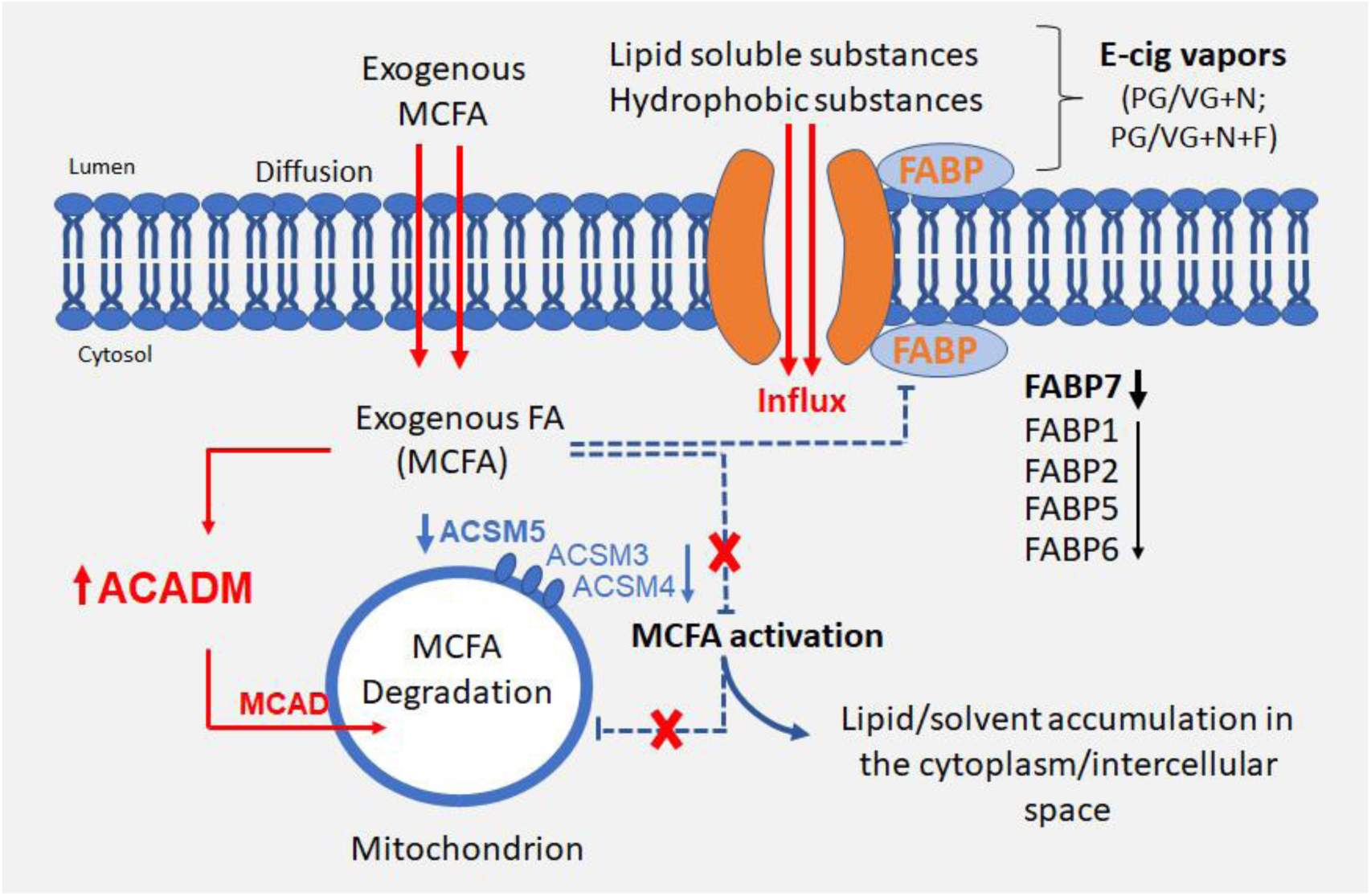
**Schematic illustration of the effect of 5% ECVE on VF epithelial lipid homeostasis**. Exogenous fatty acids, that likely contain medium-chain fatty acids/triglycerides (MCFA), diffuse across the plasma membrane into the cytosol and activate ACADM gene encoding an enzyme, medium-chain acyl -CoA dehydrogenase (MCAD) in mitochondria. Simultaneously, an increased deposition of MCFA in the cytosol inhibits synthetic reactions of MCFA with acyl-CoA and adenosine triphosphate (ATP) via downregulation of ACSM5, ASCM4 and ASCM3 in the cytosol. As a result, MCFA are not translocated into mitochondria and remain in the cytoplasm. Besides the failure in MCFA activation, ECVE also likely targets fatty acid binding proteins, FABP, that facilitate transport of lipids and/or lipid soluble substances across the plasma membrane and regulate lipid cytoplasmic content. Structural changes of the transport system due to downregulation of FABPs can cause uncontrolled influx of lipid and/or solvent particles into the cells. Lipid droplets and lipophilic solvents can also deposit between the plasma membranes in the intercellular space.

Another target of ECVE could be inactivation of fatty acid binding proteins, FABPs, in the plasma membrane that assist in the fatty acid transport across the membrane and regulate fatty acid content (23). Inactivation of these proteins can lead to uncontrolled influx of fatty acids or lipophilic solvents into the cells. So far, the role of FABPs has been mostly studied in tissues with the high lipid and/or glucose metabolism such as adipose tissue, liver, intestine, heart or skeletal muscles (33–35). Several FABP isomorphs, such as FABP7, have been also found in the epidermis, brain or lungs (23, 33), however, their role in these tissues is poorly understood. It has been shown that dysfunction of FABPs in skeletal and cardiac muscles causes lipotoxicity which leads to metabolic diseases (23). Pharmacological manipulations of FABP functions can, therefore, serve as promising future targets to correct lipid fluxes and regain metabolic homeostasis (33). Whether targeting *FABPs* can also prevent cellular damage in the stratified VF epithelium in response to ECVE remains to be investigated.

Collectively, our findings revealed two important aspects that deserve further attention. First, in line with previous studies (7, 11) there is an urgent need to investigate the physiological effects of e-liquids currently on the market, especially now, as the market is still evolving, and new products become available. We detected significant differences in expression levels of genes involved in medium fatty acid/ triglyceride metabolism in commercially available products, with nicotine and nicotine and flavor. In general, medium-chain triglycerides represent a major risk factor associated with vaping along with vitamin E acetate (6) and belong to priority toxicants to measure in bronchoalveolar lavage fluid in patients with acute respiratory illness (6). Wide testing of commercial products is necessary to identify their potential risks to the individual’s health.

Second, vaping, undoubtedly, represents a high potential risk to voice production and airway protection, in acute setting as well as prospective long-term chronic conditions. The healthy VF are strongly dependent on the compact epithelial barrier as they undergo constant collisions during the vibratory cycle. During conversational talk, average vibratory rate of the VF ranges from 100 to 150 hertz (Hz) for males, 180 to 250 Hz for females (Hz units means number of vibratory cycles per second), which further increases during singing and may go up to 1000 Hz in soprano operatic singers (36, 37). Detachment of apical cell layers burdened with lipid content in combination with constant mechanical stress can lead, ultimately, to chronic VF pathophysiological changes and life-long impairment of voice quality. Given the fact that vaping is highly popular among high school students and young adults still pursuing their professional carriers is especially alarming, as occupation-related voice disorders prevail among treatment -seeking individuals (38–40). Future clinical studies are necessary to confirm whether e-cig users are more susceptible to chronic voice disorders as compared to conventional cigarette smokers and non-smokers.

## Material and Methods

### Study design

The primary goal of this study was to determine the effect of vaporized e-liquids on VF mucosa remodeling and inflammation and test whether we can recapitulate clinical findings in in vitro experimental conditions. We first evaluated expression pattern of key structural and functional VF epithelial genes and then provided the comprehensive genomic analysis focusing on the expression of human chemokine and cytokines that are associated with VF mucosal immune responses and inflammation. The ECVE effects were measured relative to control VF mucosae treated with plain medium and we tested three different experimental conditions with PG/VG only, PG/VG and nicotine and PG/VG, nicotine and flavor – the most popular type of e-cig. We also aimed to further investigate the possible mechanisms responsible for the epithelial injury and provided comprehensive genomic analysis focusing on human fatty acid metabolism that identified medium-chain fatty acids as priority toxicants causing epithelial cell detachment. Overall, thirty-eight 3D constructs were generated in this study to provide at least two biological replicates for each experimental procedure. 3D constructs composed of vocal fold basal progenitors that did not attach to the collagen constructs properly were excluded from the study. A major limitation of this study was the absence of immune cells in our in vitro system, which could influence the transcript levels of chemokines and cytokines expressed by mucosal cells. CT values, fold change regulation values and p-Values for all tested genes are provided in Supplementary Materials. This in vitro system provided mechanistic insight into the effect of vaping on the VF mucosa health and function and serves as a necessary foundation for studying vaping in combination with infectious agents involved in acute epiglottitis and laryngitis.

### Human iPS-GFP cell culture and differentiation

For differentiation of hiPSC-derived VF epithelium we followed our recently published protocol (15). Briefly, human iPS-GFP IMR-90-4 reporter cells were maintained in an undifferentiated state in mTesr1 media on plates coated with Matrigel and were routinely passaged with Versene (StemCell Technologies, Vancouver, CA) in a ratio of 1:6. When cells reached 80% confluency, definitive endoderm induction was performed (Day 1) using RPMI medium with Glutamax (Gibco, Life Technologies) supplemented with 100ng/ml Activin A (Peprotech, Rocky Hill, NJ, USA), 25ng/ml Wnt 3a (R&D System, Minneapolis, MN, USA) and 10μM Y-27632 (R&D System, Minneapolis, MN, USA) for one day and RPMI media with Glutamax supplemented with 100ng/ml Activin A and 0.2% fetal bovine serum (FBS) for additional two days. At Day 4, anterior foregut endoderm (AFE) differentiation was performed. RPMI medium was replaced by DMEM/F12 medium with Glutamax (Gibco, Life Technologies) supplemented with N2 and B27 supplements (Gibco, Life Technologies), ascorbic acid 0.05 mg/ml (Milipore Sigma, St. Louis, MO), monothioglycerol (MTG) 0.4 mM (Milipore Sigma, St. Louis, MO) (here refer as DMEM basal medium), 200ng/ml Noggin (R&D System, Minneapolis, MN, USA) and 10μM SB431542 (Tocris, Minneapolis, MN, USA) for four days. Medium was changed daily. After 4 days of AFE treatment, at Day 8, AFE derived cells were differentiated into VF basal progenitors (VBP) to induce expression of stratified markers for an additional 4 days. We used DMEM basal medium supplemented with 1% penicillin-streptomycin (Invitrogen, Carlsbad, CA, USA) and a combination of FGF growth factors (FGF2 250ng/ml; FGF7 100ng/ml and FGF10 100ng/ml). FGFs signals were purchased from R&D System (Minneapolis, MN, USA). Medium was changed every other day. Half-way through their differentiation, at Day 10, VFB were mildly detached with 0.05% TE trypsin in EDTA (Gibco Life Technologies) and transferred to the top of the collagen-fibroblasts constructs to create organotypic VF mucosae.

### Organotypic VF mucosa cultures

One day before VBP re-seeding (Day 9), collagen constructs were prepared (15, 41) by combining high-concentration rat tail collagen (4mg/ml; 80% final volume BD Biosciences) and 10xDMEM (10% final volume; Millipore Sigma, St. Louis, MO, USA) on ice and adjusting pH (between 7.2 and 7.3) with 1N NaOH. VF primary fibroblasts 21T cells, passage P5 - P6, were resuspended in ice-cold FBS (10% final volume; 500 000 cells/ml final volume) and added to a collagen mixture. A mixture of collagen gel and VF fibroblasts was plated on transwell culture inserts (Corning, Millipore Sigma, St. Louis, MO, USA), 2ml per a 6-well culture insert, and solidified for one hour in a tissue incubator at 5% CO, 37^0^C degrees. After one hour, collagen was gently detached with a pauster pipette and constructs were flooded with DMEM basal medium, returned into an incubator and left at least 24 hours to allow for gel contraction. The next day, VBP (Day 10) were mildly trypsinized and plated on collagen constructs at high density in 100 μl DMEM basal medium supplemented with high concentration of FGF2 (250ng/ml), FGF10 (100ng/ml) and FGF7 (100ng/ml). Cells were allowed to attach for at least two hours and were then were flooded with DMEM basal medium supplemented with high levels of FGFs as mentioned above and cultivated for an additional two days to complete VBP differentiation (Day 12). On Day 12, DMEM basal medium was changed for conditional flavonoid adenine dinucleotide (FAD) medium supplemented with high levels of FGFs and VBP were further differentiated as submerged cultures for 2 days. At Day 14, conditional culture medium with FGFs was aspired from the upper inserts and cell were cultivated at the Air/Liquid interface (A/Li). The A/Li was performed in conditional FAD medium with FGFs for the first 4 days and plain FAD medium for an additional 2 weeks. FAD medium was freshly prepared every week. It consisted of the DMEM medium and F12 in ratio 1:3 (Gibco Life Technologies), supplemented with 2.5 ml FBS, 0.4 μg/ml hydrocortisone (Millipore Sigma, St. Louis, MO, USA) 8.4ng /ml cholera toxin (Millipore Sigma, St. Louis, MO, USA), 5 μg/ml insulin (Millipore Sigma, St. Louis, MO, USA), 24μg/ml adenine (Millipore Sigma, St. Louis, MO, USA), 10ng/ml epidermal growth factor, 1% penicillin-streptomycin (Invitrogen, Carlsbad, CA, USA). In submerged cultures, 1 ml of FAD was applied on transwells with collagen constructs and 2 ml were applied in the basolateral chamber. FAD was changed every other day. To create the A/Li, FAD medium was placed in the basolateral chamber only and changed three times a week. Conditional FAD medium was formed by cultivation of FAD with human primary VFF 21T cells for 24hours in 37^0^C in 5% CO2-humidified atmosphere. After 24hours, medium was collected, sterile-filtered and stored at -20^0^ C. The ratio of 30:70 (30% for conditional and 70% for fresh FAD medium) was used in the experiment.

### Preparation of the electronic cigarette vapor extract (ECVE)

ECVE (100%) was generated as recently described (15). Briefly, e-cigarette vapors (sold by Infinite Vapor, infinitevapor.com) were bubbled through 30 ml the DMEM/F12 medium (Gibco Technologies) in a disposable 50 ml tube with the use of an experimenter-operated syringe. Human vaping was modeled with two short puffs (2 seconds) with long delays between puffs (30 seconds).Two short puffs were repeated 15x (30 puffs total), which was equivalent to three conventional cigarettes used in our previous study (15) for each experimental condition. Aerosolized vapors that were drawn through the end of the e-cigarette during vaping, were bubbled through the DMEM medium. The obtained medium was considered 100% EVCE. To ensure standardization between experiments, ECVE was sterile-filtered through a 0.2 mm filter, aliquoted, and stored at -80^0^C. Before usage, ECVE was quickly thawed and diluted with the FAD medium to the indicated concentration and used the same day. We prepared three different 100% ECVE extracts using three distinct e-liquid cartridges for three different experimental conditions: PG/VG only (vehicle control), 0% nicotine (Hell Vapors; batch number: G0391); the e-liquid with PG/VG with 1.8% nicotine (Hells Vapors, batch number: C1291) and the e-liquid with PG/VG with 1.2% nicotine and Unicorn Poop flavor (Drip Star, batch number: E0891).

### Functional experiments using 5% EVCE

To mimic chronic exposure of VF mucosae to ECVE, inserts (upper chamber) containing engineered VF mucosae at Day 32 of differentiation were flooded with FAD medium supplemented with 5% ECVE for 1 week. We used 5% ECVE with PG/VG only, 5% ECVE with PG/VG+N and 5% ECVE with PG/VG+N+F. The lower chamber was flooded with plain FAD medium. Engineered VF mucosae flooded with plain FAD medium were used as negative controls. Medium was changed every day in both chambers. After one week of exposure to the 5% ECVE engineered VF mucosae were characterized with immunohistochemistry (IHC) and quantitative polymerase chain reaction (qPCR) to investigate expression levels of clinically relevant genes in VF mucosal cells. We also performed Oil Red O stain on frozen VF mucosa sections to detect lipid particles and TEM to evaluate the ultrastructure of VF apical epithelial cell layers. Thirty-eight 3D constructs were generated in this study to enable at least two biological replicates for each experimental procedure. VBP that did not attach to the collagen constructs properly were excluded from the experiment.

### Cryosectioning and Oil Red O stain

At Day 39, medium was aspired from the control and 5% ECVE exposed VF mucosae; 3D constructs were briefly washed in PBS immediately embedded in separate molds with O.C.T (Sakura, Hayward, CA) mounting medium and placed on dry ice to freeze tissues. Blocks were stored in the freezer - 80^0^C. Before cryosectioning, blocks were removed from the freezer, allowed to warm up to -21^0^C in a cryostat (Leica CM3050S) and cut to 5 μm thick sections, (chamber temperature -21^0^C and objective temperature -21^0^C). Sections were collected on pre-coated slides and immediately used for Oil Red Oil staining. Oil Red staining was performed using Oil Red O Stain Kit (Lipid Stain) (Abcam, Cambridge, CA, USA) following manufacturer’s protocol. Samples were contra-stained with hematoxylin and mounted with Vectashield without DAPI (Vector Laboratories; Peterborough, UK). Images were taken with a Nikon Eclipse E600 with a camera Olympus DP71, and were adjusted for brightness using the installed DP 71 software, Olympus Corporation.

### Immunohistochemistry (IHC) of collagen gel constructs

At Day 39, collagen gel constructs were first washed in PBS, fixed in fresh 4% paraformaldehyde for 15 min at RT and embedded in histogel (Thermo Fisher Scientific, Waltham, MA, USA). Constructs were dehydrated in a series of ethanol, treated with xylene, embedded in paraffin, and cut to 5μm thick serial sections. Sections were then deparaffinized, rehydrated and stained using a standard IHC protocol (15). Antigen retrieval was performed by heating sections in sodium citrate pH = 6 at 80^0^C water bath for two hours. Primary antibodies used included rabbit anti-Laminin alfa 5 diluted at 1:100 (Abcam; Cambridge, UK), anti-rabbit cytokeratin K14 diluted to 1:250 (Proteintech, Rosemont, IL, USA), anti-rabbit cytokeratin K13 diluted to 1:100 (Abcam, Cambridge, CA, USA), anti-rabbit E-Cadherin diluted to 1:250 (Cell Signaling, Danvers, MA, USA), anti-mouse p63 diluted to 1:100 (Santa Cruz Biotechnology, Dallas, TX, USA), and anti-mouse MUC1 and MUC4 diluted to 1:200 (both Abcam, Cambridge, CA, USA). Secondary antibodies used were Alexa Fluor TM 488 goat anti-rabbit (Invitrogen, Carlsbad, CA, USA) at the dilution 1:500, Cy3-cojugated goat anti-mouse at the dilution 1:200 (both Jackson ImmunoResearch; West Grove, PA, USA). They were applied 1h and 30 min at RT. Slides were mounted using Vectashield with DAPI (Vector Laboratories; Peterborough, UK). Images were taken with a Nikon Eclipse E600 with a camera Olympus DP71, and were adjusted for brightness using the installed DP 71 software, Olympus Corporation.

### Transmission electron microscopy

For transmission electron microscopy, gels were fixed in 2.5% glutaraldehyde and 2% paraformaldehyde in a 0.1M sodium cacodylate buffer (pH 7.4) overnight at 4^0^C and processed using routine techniques. Briefly, gels were washed in a 0.1M sodium cacodylate buffer and postfixed in 1% osmium tetroxide in the same buffer for 2 h at room temperature. Tissues were dehydrated in graded ethanol series, rinsed twice in propylene oxide, and embedded in Epon 812 epoxy resin (Polysciences, Inc., Warrington, PA) under vacuum. Finally, the samples were flat embedded between glass slides. After resin polymerization, one of the two glass slides were removed, and blank resin cylinders were glued to the sections. The gels were thin sectioned for TEM using a Leica EM UC6 Ultramicrotome and stained with Reynolds lead citrate and 8% uranyl acetate in 50% EtOH to increase contrast. The sections were viewed with a Philips CM120 electron microscope and images were captured with a MegaView III side mounted digital camera.

### Isolation of VF mucosal cells for qPCR

To isolate populations of VF mucosal cells from the whole constructs, collagen gel was first dissolved using collagenase (Gibco™ Collagenase, Type I, Powder; 17018029 Gibco™). Briefly, old medium was aspired, and constructs were washed twice in PBS. Collagenase type I at working concentration 100U/ml was added to the upper (1ml) and lower chambers (2ml), with VF mucosa being fully submerged. Constructs were incubated at 37^0^C for at least 2 – 3 hours, or until the collagen completely dissolved and cells got loose. The cell suspension was then transferred into a 15ml conical tube. Cells were centrifuged for 5 min at 10,000 rpm, washed and resuspended in PBS and transferred into a 1.5ml tube. Cells were again centrifuged for 5 min at 10,000 rpm, the supernatant was aspired, and cell pellet was stored at -80^0^C.

### RNA isolation and qPCR

Cells isolated from the whole constructs were used for RNA isolation using ReliaPrep^TM^ RNACell Miniprep System (Promega, Madison, WI) following manufacturer’s protocol. One thousand ng of RNA was reverse transcribed to cDNA using reverse transcription reagents (Go Script, Promega, Madison, WI) per manufacturer’s protocol. Total volume of 0.4μl of cDNA was used per 20μl real time qPCR reaction using Power Up Sybr Green Master Mix (Applied Biosystem, Foster City, CA, USA) and run for 40 cycles in triplicates on a 7500 Fast Real Time PCR System machine (Applied Biosystem, Foster City, CA, USA), according to manufacturer’s instructions. Gene specific primers are listed in Table 1. Relative gene expression, normalized to beta-Actin (Delta CT), and control VF mucosae (Delta Delta CT), was calculated as fold change using the 2(-Delta Delta CT) method. If undetected, a cycle number 40 was assigned to allow fold change calculations. Data are presented as the average of the two biological and three technical replicates ± standard error of the mean. One-Way ANOVA of Variance for Independent or Correlated Samples analysis along with Tukey HSD test were used to confirm statistical significance in gene expression {p ≤ 0.05 (*) and p ≤ 0.01 (**)].

**Table 1:**
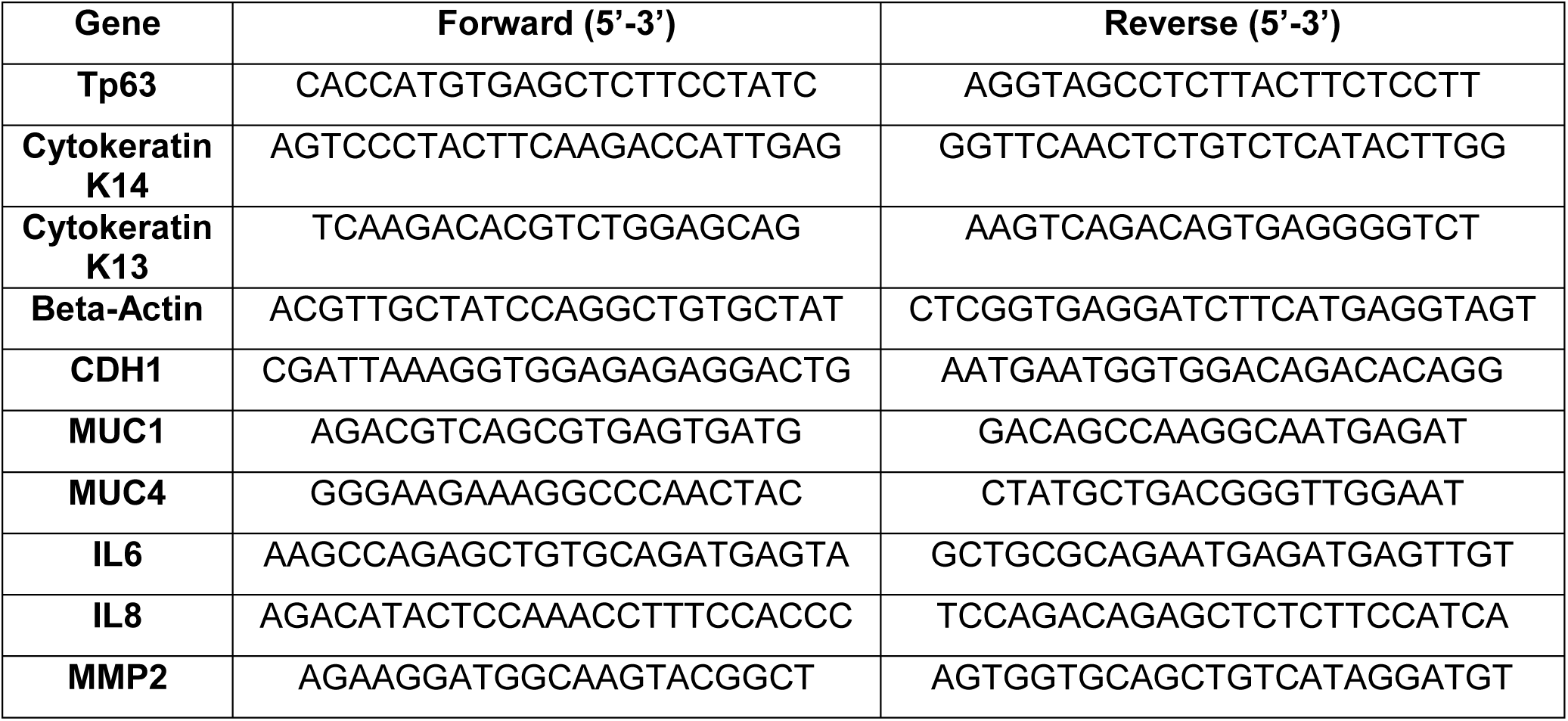
Genes and primers used in the stud

### RT2 PCR profiling

Cells isolated from whole constructs were used for RNA isolation using ReliaPrep^TM^ RNACell Miniprep System (Promega, Madison, WI) following manufacturer’s protocol, Five hundred ng of RNA was reverse transcribed to cDNA using a reverse transcription RT^2^ First Strand Kit (Qiagen, Hilden, Germany) that is recommended to use in combination with RT^2^ Profiler PCR Arrays and we followed manufacturer’s protocol. Immediately after cDNA reverse transcription we followed with RT^2^ Profiler PCR Arrays. The cDNA was first diluted with nuclease-free water and then added to the RT^2^ SYBR Green ROX^TM^ qPCR Mastermix (Qiagen, Hilden, Germany) according to manufacturer’s protocol. We used RT^2^ Profiler PCR Arrays for Human Cytokines and Chemokines (PAHS-150Z), and Fatty Acid Metabolism (PAHS-007Z), both purchased from Qiagen (Qiagen, Hilden, Germany). A RT2 Profiler PCR Array Format E 384 (4 x 96) was used to be able to screen 4 different samples on one plate. The 384 (4 x 96) option contained 4 replicate primer assay for each 84 pathway-focused genes and 5 replicate primer assay for each housekeeping (reference) gene; beta-Actin (*ACTB*), beta-2-Microglobulin (*B2M*), Glyceraldehyde-3-phosphate dehydrogenase (*GAPDH*), Hypoxanthine phosphoribosyltransferase 1 (*HPRT1*), and Ribosomal protein, large, P0 (*RPLP0*). Assays for 5 housekeeping genes enabled normalization of data. Control and each test group were run in a triplicate (n=3, 12 samples total). We used a real-time cycler Quant Studio 5 Applied Biosystems (Foster City, CA, USA). Cycling conditions were: 95.0°C for 10:00 minutes for Stage 1; Stage 2, 95.0°C for 15 seconds followed by 60.0°C for 1:00 minute with 40 repeats (40 cycles) and the Dissociation Stage 3. CT values were exported to an Excel file to create a table of CT values (Supplemental Tables S2 and S5). These tables were then uploaded on to the data analysis web portal at http://www.qiagen.com/geneglobe. Samples were assigned to controls (Control) and test groups: PG/VG (Group 1), PG/VG+N (Group 2), PG/VG+N+F (Group 3). The CT cut-off was set to 35. CT values were normalized based on a Manual Selection of reference (housekeeping) genes. The data analysis web portal calculated fold change/regulation using delta delta CT method, in which delta CT was calculated between gene of interest (GOI) and an average of reference genes (HKG), followed by delta-delta CT calculations (delta CT (Test Group)-delta CT (Control Group)). Fold change was then calculated using 2∧ (-delta delta CT) formula. Fold-Regulation represents fold-change results in a biologically meaningful way. Fold-change values greater than one indicate a positive- or an up-regulation, and the fold-regulation was equal to the fold-change. Fold-change values less than one indicate a negative or down-regulation, and the fold-regulation was the negative inverse of the fold-change (Supplemental Tables S3 and S6). The fold-change threshold was set to 2. The p-values were calculated based on a Student’s t-test of the replicate 2∧(-Delta Delta CT) values for each gene in the control group and treatment groups, and p-values less than 0.05 were considered as significant. The p-value calculation used was based on parametric, unpaired, two-sample equal variance, two-tailed distribution.

### Statistical analysis

One-Way ANOVA of Variance for Independent or Correlated Samples analysis along with Tukey HSD test were used to confirm statistical significance in gene expression [p ≤ 0.05 (*) and p ≤ 0.01 (**)] in structural and functional epithelial genes. For Human Cytokines and Chemokine expression and Fatty Acid metabolism expression, the p-values were calculated based on a Student’s t-test of the replicate 2∧(-Delta Delta CT) values for each gene in the control group and treatment groups, and p-values less than 0.05 were considered as significant. The p-value calculation used was based on parametric, unpaired, two-sample equal variance, two-tailed distribution.

## Author contributions

V.L. and S.L.T. designed research; V.L. performed experiments; S.L.T supervised the work; V.L and S.L.T wrote the manuscript. Authors approved the final manuscript.

## Ethical statement

All stem cell work in this investigation was approved by the Stem Cell Research Oversight Committee at the University of Wisconsin Madison (SC-2015-0008).

## Acknowledgement

This work was funded the grants NIH-NIDCD R01 04336 and R01 012773. We gratefully acknowledge Sierra Raglin for her expert assistance with the construct sample preparation for this study. We also gratefully acknowledge Stephanie Bartley for her assistance in ECVE preparation.

---

**Supplemental Table S1:** Table of genes analyzed in the RT_2_ Profiler PCR Arrays --Human Cytokine and Chemokines (PAHS-150Z)

| Position | RefSeq Number | Symbol | Description |
| --- | --- | --- | --- |
| A01 | NM_004797 | **ADIPOQ** | Adiponectin, C1Q and collagen domain containing |
| A02 | NM_001200 | **BMP2** | Bone morphogenetic protein 2 |
| A03 | NM_130851 | **BMP4** | Bone morphogenetic protein 4 |
| A04 | NM_001718 | **BMP6** | Bone morphogenetic protein 6 |
| A05 | NM_001719 | **BMP7** | Bone morphogenetic protein 7 |
| A06 | NM_001735 | **C5** | Complement component 5 |
| A07 | NM_002981 | **CCL1** | Chemokine (C-C motif) ligand 1 |
| A08 | NM_002986 | **CCL11** | Chemokine (C-C motif) ligand 11 |
| A09 | NM_005408 | **CCL13** | Chemokine (C-C motif) ligand 13 |
| A10 | NM_002987 | **CCL17** | Chemokine (C-C motif) ligand 17 |
| A11 | NM_002988 | **CCL18** | Chemokine (C-C motif) ligand 18 (pulmonary and activation-regulated) |
| A12 | NM_006274 | **CCL19** | Chemokine (C-C motif) ligand 19 |
| B01 | NM_002982 | **CCL2** | Chemokine (C-C motif) ligand 2 |
| B02 | NM_004591 | **CCL20** | Chemokine (C-C motif) ligand 20 |
| B03 | NM_002989 | **CCL21** | Chemokine (C-C motif) ligand 21 |
| B04 | NM_002990 | **CCL22** | Chemokine (C-C motif) ligand 22 |
| B05 | NM_002991 | **CCL24** | Chemokine (C-C motif) ligand 24 |
| B06 | NM_002983 | **CCL3** | Chemokine (C-C motif) ligand 3 |
| B07 | NM_002985 | **CCL5** | Chemokine (C-C motif) ligand 5 |
| B08 | NM_006273 | **CCL7** | Chemokine (C-C motif) ligand 7 |
| B09 | NM_005623 | **CCL8** | Chemokine (C-C motif) ligand 8 |
| B10 | NM_000074 | **CD40LG** | CD40 ligand |
| B11 | NM_000614 | **CNTF** | Ciliary neurotrophic factor |
| B12 | NM_000757 | **CSF1** | Colony stimulating factor 1 (macrophage) |
| C01 | NM_000758 | **CSF2** | Colony stimulating factor 2 (granulocyte-macrophage) |
| C02 | NM_000759 | **CSF3** | Colony stimulating factor 3 (granulocyte) |
| C03 | NM_002996 | **CX3CL1** | Chemokine (C-X3-C motif) ligand 1 |
| C04 | NM_001511 | **CXCL1** | Chemokine (C-X-C motif) ligand 1 (melanoma growth stimulating activity, alpha) |
| C05 | NM_001565 | **CXCL10** | Chemokine (C-X-C motif) ligand 10 |
| C06 | NM_005409 | **CXCL11** | Chemokine (C-X-C motif) ligand 11 |
| C07 | NM_000609 | **CXCL12** | Chemokine (C-X-C motif) ligand 12 |
| C08 | NM_006419 | **CXCL13** | Chemokine (C-X-C motif) ligand 13 |
| C09 | NM_022059 | **CXCL16** | Chemokine (C-X-C motif) ligand 16 |
| C10 | NM_002089 | **CXCL2** | Chemokine (C-X-C motif) ligand 2 |
| C11 | NM_002994 | **CXCL5** | Chemokine (C-X-C motif) ligand 5 |
| C12 | NM_002416 | **CXCL9** | Chemokine (C-X-C motif) ligand 9 |
| D01 | NM_000639 | **FASLG** | Fas ligand (TNF superfamily, member 6) |
| D02 | NM_000175 | **GPI** | Glucose-6-phosphate isomerase |
| D03 | NM_000605 | **IFNA2** | Interferon, alpha 2 |
| D04 | NM_000619 | **IFNG** | Interferon, gamma |
| D05 | NM_000572 | **IL10** | Interleukin 10 |
| D06 | NM_000641 | **IL11** | Interleukin 11 |
| D07 | NM_000882 | **IL12A** | Interleukin 12A (natural killer cell stimulatory factor 1, cytotoxic lymphocyte maturation factor 1, p35) |
| D08 | NM_002187 | **IL12B** | Interleukin 12B (natural killer cell stimulatory factor 2, cytotoxic lymphocyte maturation factor 2, p40) |
| D09 | NM_002188 | **IL13** | Interleukin 13 |
| D10 | NM_000585 | **IL15** | Interleukin 15 |
| D11 | NM_004513 | **IL16** | Interleukin 16 |
| D12 | NM_002190 | **IL17A** | Interleukin 17A |
| E01 | NM_052872 | **IL17F** | Interleukin 17F |
| E02 | NM_001562 | **IL18** | Interleukin 18 (interferon-gamma-inducing factor) |
| E03 | NM_000575 | **IL1A** | Interleukin 1, alpha |
| E04 | NM_000576 | **IL1B** | Interleukin 1, beta |
| E05 | NM_000577 | **IL1RN** | Interleukin 1 receptor antagonist |
| E06 | NM_000586 | **IL2** | Interleukin 2 |
| E07 | NM_021803 | **IL21** | Interleukin 21 |
| E08 | NM_020525 | **IL22** | Interleukin 22 |
| E09 | NM_016584 | **IL23A** | Interleukin 23, alpha subunit p19 |
| E10 | NM_006850 | **IL24** | Interleukin 24 |
| E11 | NM_145659 | **IL27** | Interleukin 27 |
| E12 | NM_000588 | **IL3** | Interleukin 3 (colony-stimulating factor, multiple) |
| F01 | NM_000589 | **IL4** | Interleukin 4 |
| F02 | NM_000879 | **IL5** | Interleukin 5 (colony-stimulating factor, eosinophil) |
| F03 | NM_000600 | **IL6** | Interleukin 6 (interferon, beta 2) |
| F04 | NM_000880 | **IL7** | Interleukin 7 |
| F05 | NM_000584 | **CXCL8** | Interleukin 8 |
| F06 | NM_000590 | **IL9** | Interleukin 9 |
| F07 | NM_002309 | **LIF** | Leukemia inhibitory factor (cholinergic differentiation factor) |
| F08 | NM_000595 | **LTA** | Lymphotoxin alpha (TNF superfamily, member 1) |
| F09 | NM_002341 | **LTB** | Lymphotoxin beta (TNF superfamily, member 3) |
| F10 | NM_002415 | **MIF** | Macrophage migration inhibitory factor (glycosylation-inhibiting factor) |
| F11 | NM_005259 | **MSTN** | Myostatin |
| F12 | NM_018055 | **NODAL** | Nodal homolog (mouse) |
| G01 | NM_020530 | **OSM** | Oncostatin M |
| G02 | NM_002704 | **PPBP** | Pro-platelet basic protein (chemokine (C-X-C motif) ligand 7) |
| G03 | NM_000582 | **SPP1** | Secreted phosphoprotein 1 |
| G04 | NM_003238 | **TGFB2** | Transforming growth factor, beta 2 |
| G05 | NM_000460 | **THPO** | Thrombopoietin |
| G06 | NM_000594 | **TNF** | Tumor necrosis factor |
| G07 | NM_002546 | **TNFRSF11B** | Tumor necrosis factor receptor superfamily, member 11b |
| G08 | NM_003810 | **TNFSF10** | Tumor necrosis factor (ligand) superfamily, member 10 |
| G09 | NM_003701 | **TNFSF11** | Tumor necrosis factor (ligand) superfamily, member 11 |
| G10 | NM_006573 | **TNFSF13B** | Tumor necrosis factor (ligand) superfamily, member 13b |
| G11 | NM_003376 | **VEGFA** | Vascular endothelial growth factor A |
| G12 | NM_002995 | **XCL1** | Chemokine (C motif) ligand 1 |
| H01 | NM_001101 | **ACTB** | Actin, beta |
| H02 | NM_004048 | **B2M** | Beta-2-microglobulin |
| H03 | NM_002046 | **GAPDH** | Glyceraldehyde-3-phosphate dehydrogenase |
| H04 | NM_000194 | **HPRT1** | Hypoxanthine phosphoribosyltransferase 1 |
| H05 | NM_001002 | **RPLP0** | Ribosomal protein, large, P0 |
| H06 | SA_00105 | **HGDC** | Human Genomic DNA Contamination |
| H07 | SA_00104 | **RTC** | Reverse Transcription Control |
| H08 | SA_00104 | **RTC** | Reverse Transcription Control |
| H09 | SA_00104 | **RTC** | Reverse Transcription Control |
| H10 | SA_00103 | **PPC** | Positive PCR Control |
| H11 | SA_00103 | **PPC** | Positive PCR Control |
| H12 | SA_00103 | **PPC** | Positive PCR Control |

**Supplemental Table S2:**
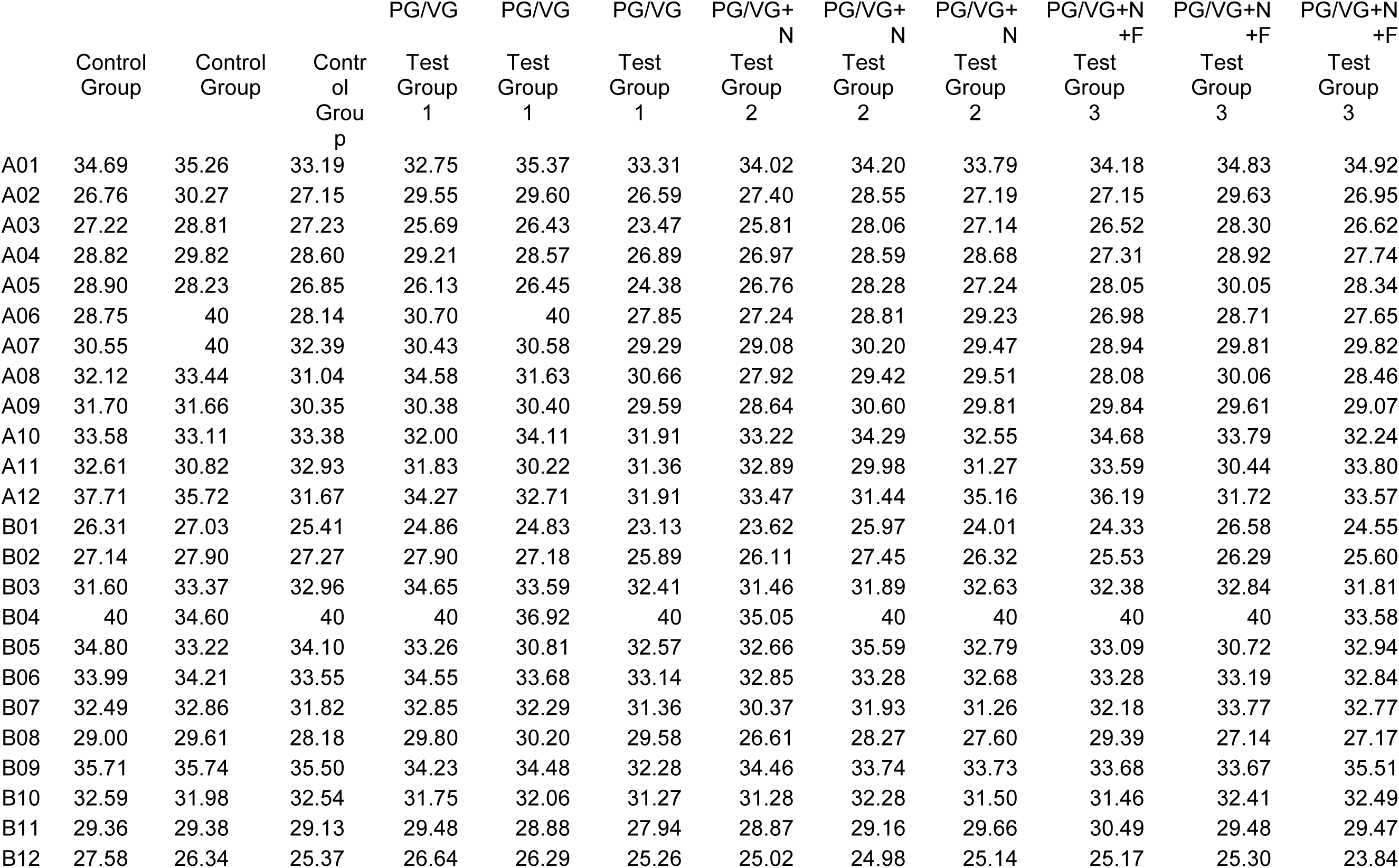
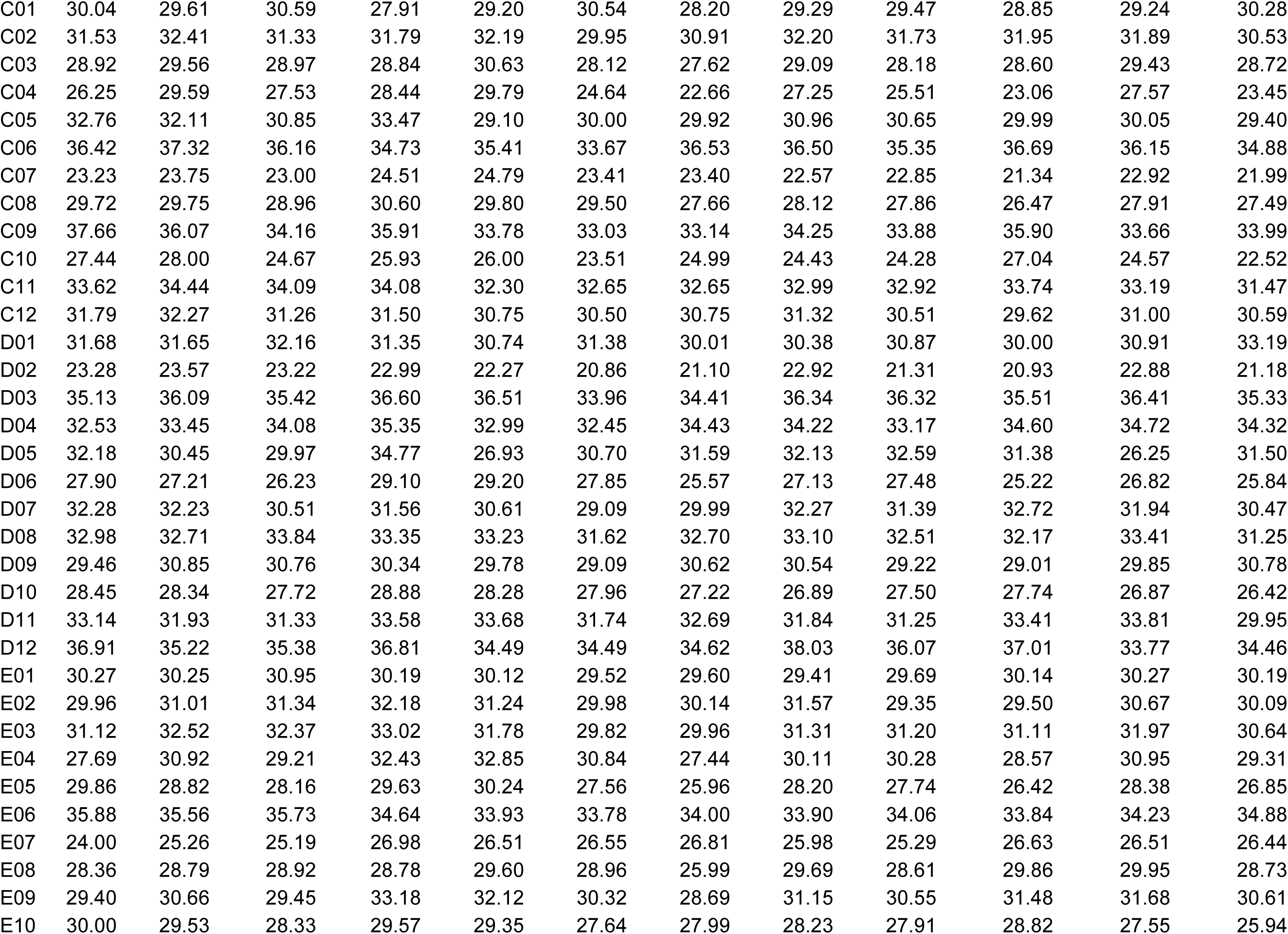
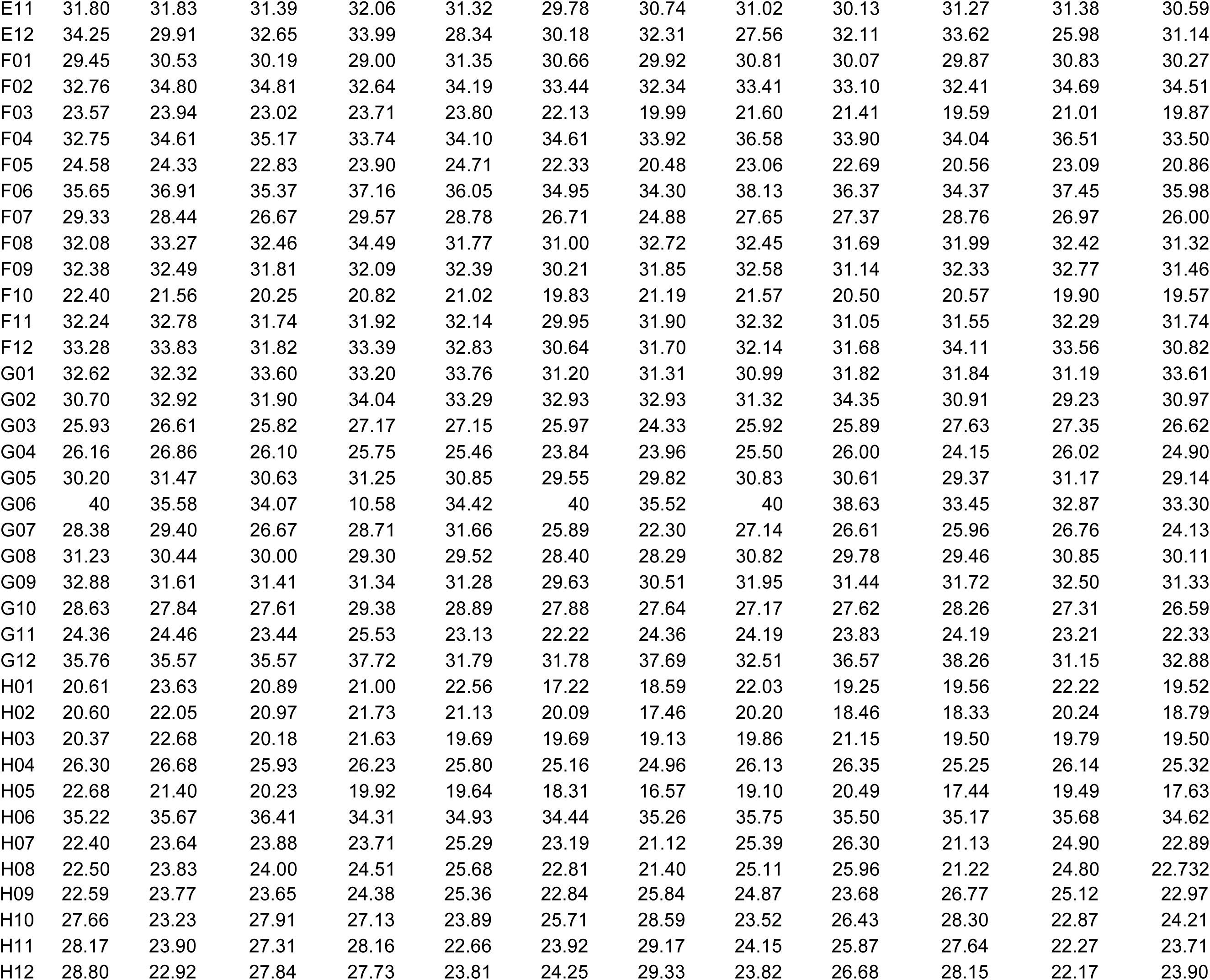
List of CT values of human cytokines and chemokines uploaded on to the data analysis web portal at http://www.qiagen.com/geneglobe

**Supplemental Table S3:** Fold regulation and p-values of genes associated mucosal inflammation. Fold regulation is compared to controls

| Position | Gene Symbol | Group 1 |  | Group 2 |  | Group 3 |  |
| --- | --- | --- | --- | --- | --- | --- | --- |
|  |  | Fold Regulation | p-Value | Fold Regulation | p-Value | Fold Regulation | p-Value |
| A01 | **ADIPOQ** | -1.34 | 0.949612 | -2.65 | 0.07756 | -4.34 | 0.022261 |
| A02 | **BMP2** | -2.93 | 0.162542 | -2.56 | 0.214097 | -3.07 | 0.150726 |
| A03 | **BMP4** | 2.89 | 0.048146 | -1.93 | 0.019491 | -2.22 | 0.007207 |
| A04 | **BMP6** | -1.13 | 0.333216 | -1.62 | 0.024908 | -1.6 | 0.015384 |
| A05 | **BMP7** | 2.49 | 0.044874 | -2.19 | 0.15756 | -5.99 | 0.051444 |
| A06 | **C5** | -2.99 | 0.26999 | 1.42 | 0.521938 | 2.12 | 0.946686 |
| A07 | **CCL1** | 2.87 | 0.296137 | 2.57 | 0.391479 | 2.56 | 0.40323 |
| A08 | **CCL11** | -2.17 | 0.211494 | 2.93 | 0.009991 | 2.97 | 0.003055 |
| A09 | **CCL13** | 1.06 | 0.855139 | -1.1 | 0.634625 | -1 | 0.89173 |
| A10 | **CCL17** | -1.27 | 0.847791 | -3.24 | 0.228373 | -3.95 | 0.188997 |
| A11 | **CCL18** | -1.03 | 0.717673 | -1.94 | 0.633077 | -4.78 | 0.471747 |
| A12 | **CCL19** | -1.08 | 0.653829 | -2.16 | 0.832994 | -2.47 | 0.749384 |
| B01 | **CCL2** | 1.93 | 0.005131 | 1.01 | 0.765233 | -1.59 | 0.049993 |
| B02 | **CCL20** | -1.49 | 0.161355 | -1.85 | 0.180155 | -1.1 | 0.733896 |
| B03 | **CCL21** | -3.81 | 0.076845 | -2.06 | 0.224412 | -2.76 | 0.11197 |
| B04 | **CCL22** | -2.23 | 0.29771 | -3.56 | 0.192872 | -2.69 | 0.229169 |
| B05 | **CCL24** | 1.74 | 0.560675 | -2.2 | 0.38765 | 1.02 | 0.733473 |
| B06 | **CCL3** | -1.87 | 0.158638 | -1.65 | 0.368825 | -1.93 | 0.216966 |
| B07 | **CCL5** | -1.75 | 0.090679 | -1.41 | 0.254146 | -4.87 | 0.010946 |
| B08 | **CCL7** | -3.88 | 0.008 | -1.2 | 0.425252 | -1.66 | 0.946204 |
| B09 | **CCL8** | 1.24 | 0.778056 | -1.6 | 0.610014 | -1.85 | 0.510117 |
| B10 | **CD40LG** | -1.28 | 0.57854 | -2.02 | 0.329164 | -2.86 | 0.238628 |
| B11 | **CNTF** | -1.41 | 0.364815 | -3.11 | 0.121358 | -4.88 | 0.090452 |
| B12 | **CSF1** | -1.57 | 0.295875 | -1.24 | 0.661535 | -1.08 | 0.72296 |
| C01 | **CSF2** | -1.12 | 0.769752 | -1.52 | 0.413178 | -2.21 | 0.358839 |
| C02 | **CSF3** | -1.5 | 0.0961 | -2.94 | 0.007822 | -2.76 | 0.021024 |
| C03 | **CX3CL1** | -2.11 | 0.219691 | -1.8 | 0.153513 | -2.89 | 0.038237 |
| C04 | **CXCL1** | -1.82 | 0.646625 | 1.93 | 0.39189 | 2.51 | 0.224539 |
| C05 | **CXCL10** | 1.02 | 0.615737 | -1.23 | 0.522227 | 1.25 | 0.685628 |
| C06 | **CXCL11** | -1.41 | 0.413821 | -3.25 | 0.14523 | -3.31 | 0.134927 |
| C07 | **CXCL12** | -3.83 | 0.02162 | -2.49 | 0.18512 | -1.44 | 0.200292 |
| C08 | **CXCL13** | -2.86 | 0.074479 | -1.07 | 0.980486 | 1.34 | 0.426084 |
| C09 | **CXCL16** | -1.18 | 0.626293 | -1.67 | 0.221699 | -2.4 | 0.319296 |
| C10 | **CXCL2** | 1.44 | 0.761869 | 1.35 | 0.610594 | 1.17 | 0.492224 |
| C11 | **CXCL5** | 1.01 | 0.81883 | -1.42 | 0.487189 | -1.43 | 0.658705 |
| C12 | **CXCL9** | -1.13 | 0.838286 | -1.72 | 0.288454 | -1.31 | 0.369544 |
| D01 | **FASLG** | -1.28 | 0.71466 | -1.22 | 0.660594 | -2.46 | 0.386366 |
| D02 | **GPI** | 1.22 | 0.645905 | -1.08 | 0.946249 | -1.05 | 0.721945 |
| D03 | **IFNA2** | -1.6 | 0.316888 | -2.84 | 0.149693 | -3.4 | 0.133243 |
| D04 | **IFNG** | -2.22 | 0.203707 | -4.87 | 0.106166 | -7.76 | 0.068464 |
| D05 | **IL10** | -1.95 | 0.562303 | -7.67 | 0.137885 | -1.53 | 0.501363 |
| D06 | **IL11** | -6.18 | 0.062115 | -2.48 | 0.13037 | -1.53 | 0.280914 |
| D07 | **IL12A** | 1.17 | 0.723315 | -2.36 | 0.071131 | -3.48 | 0.071281 |
| D08 | **IL12B** | -1.5 | 0.380721 | -2.45 | 0.284113 | -1.82 | 0.332568 |
| D09 | **IL13** | -1.33 | 0.399013 | -2.76 | 0.212357 | -2.44 | 0.181454 |
| D10 | **IL15** | -2.34 | 0.136928 | -1.66 | 0.493474 | -1.52 | 0.603639 |
| D11 | **IL16** | -3.72 | 0.139453 | -2.81 | 0.272895 | -4.06 | 0.244233 |
| D12 | **IL17A** | -1.61 | 0.392563 | -2.97 | 0.147803 | -2.26 | 0.414697 |
| E01 | **IL17F** | -1.4 | 0.462483 | -1.71 | 0.43161 | -2.77 | 0.209698 |
| E02 | **IL18** | -2.62 | 0.111179 | -2.43 | 0.273233 | -2.11 | 0.153393 |
| E03 | **IL1A** | -1.47 | 0.328236 | -1.43 | 0.303056 | -2 | 0.144028 |
| E04 | **IL1B** | -13.84 | 0.135316 | -3.26 | 0.214877 | -4.29 | 0.184107 |
| E05 | **IL1RN** | -2.34 | 0.19862 | -1.04 | 0.648971 | -1.03 | 0.673274 |
| E06 | **IL2** | -1.11 | 0.78615 | -1.61 | 0.473853 | -2.12 | 0.285518 |
| E07 | **IL21** | -7.39 | 0.041969 | -7.5 | 0.050909 | -11.12 | 0.035929 |
| E08 | **IL22** | -2.73 | 0.169701 | -2.15 | 0.199245 | -6.02 | 0.062789 |
| E09 | **IL23A** | -8.35 | 0.001218 | -3.98 | 0.002164 | -9.1 | 0.001272 |
| E10 | **IL24** | -1.51 | 0.238463 | -1.37 | 0.578921 | 1.06 | 0.614469 |
| E11 | **IL27** | -1.33 | 0.380075 | -1.57 | 0.533096 | -2.25 | 0.15919 |
| E12 | **IL3** | 1.33 | 0.95802 | -1.06 | 0.815109 | 1.2 | 0.484133 |
| F01 | **IL4** | -2.48 | 0.566347 | -3.77 | 0.046445 | -4.1 | 0.035901 |
| F02 | IL5 | -1.26 | 0.866017 | -1.44 | 0.388842 | -2.85 | 0.189438 |
| F03 | IL6 | -1.66 | 0.108928 | 1.75 | 0.049729 | 3.01 | 0.001628 |
| F04 | IL7 | -2.08 | 0.337616 | -3.61 | 0.146737 | -3.55 | 0.134236 |
| F05 | CXCL8 | -1.69 | 0.20717 | 1.1 | 0.942692 | 1.57 | 0.262679 |
| F06 | IL9 | -2.01 | 0.281961 | -2.77 | 0.1509 | -2.94 | 0.14391 |
| F07 | LIF | -2.35 | 0.144571 | -1.14 | 0.521211 | -1.82 | 0.501239 |
| F08 | LTA | -1.79 | 0.352771 | -2.61 | 0.128913 | -2.1 | 0.06198 |
| F09 | LTB | -1.28 | 0.403547 | -2.51 | 0.184707 | -3.31 | 0.055584 |
| F10 | MIF | -1.13 | 0.592163 | -2.6 | 0.184788 | -1.3 | 0.683908 |
| F11 | MSTN | -1.08 | 0.686789 | -2.3 | 0.180819 | -2.59 | 0.024663 |
| F12 | NODAL | -1.26 | 0.392322 | -1.47 | 0.451274 | -3.06 | 0.1661 |
| G01 | OSM | -1.87 | 0.323128 | -1.17 | 0.740635 | -2.19 | 0.552686 |
| G02 | PPBP | -6.1 | 0.0713 | -6.65 | 0.144067 | -1.23 | 0.78631 |
| G03 | SPP1 | -3.18 | 0.019935 | -1.94 | 0.054915 | -7.17 | 0.009913 |
| G04 | TGFB2 | 1.26 | 0.291319 | -1.39 | 0.263056 | -1.33 | 0.261376 |
| G05 | THPO | -1.75 | 0.070771 | -2.54 | 0.037897 | -1.85 | 0.06717 |
| G06 | TNF | 127.49 | 0.373901 | -4.02 | 0.050591 | -1.22 | 0.969815 |
| G07 | TNFRSF11B | -3.09 | 0.268953 | 2.14 | 0.359668 | 1.7 | 0.344565 |
| G08 | TNFSF10 | 1.37 | 0.659453 | -1.71 | 0.292383 | -2.54 | 0.176298 |
| G09 | TNFSF11 | 1.14 | 0.847909 | -2.05 | 0.280084 | -3.14 | 0.196984 |
| G10 | TNFSF13B | -3.29 | 0.158789 | -2.22 | 0.331585 | -2.18 | 0.332172 |
| G11 | VEGFA | -1.48 | 0.522086 | -3.33 | 0.065147 | -1.89 | 0.334883 |
| G12 | XCL1 | 2.17 | 0.328343 | -1.83 | 0.947324 | 1.17 | 0.480846 |
| H01 | ACTB | 1.34 | 0.498357 | 1.04 | 0.823407 | -1.4 | 0.424168 |
| H02 | B2M | -1.74 | 0.069622 | 1.75 | 0.198236 | 1.25 | 0.304749 |
| H03 | GAPDH | -1.22 | 0.842034 | -1.59 | 0.393738 | -1.22 | 0.668975 |
| H04 | HPRT1 | -1.37 | 0.36067 | -2.31 | 0.067216 | -2.04 | 0.08229 |
| H05 | RPLP0 | 2.17 | 0.136325 | 2.02 | 0.292426 | 2.79 | 0.046283 |

**Supplemental Table S4:** Table of genes involved in the Fatty Acid Metabolism (PAHS-007Z)

| Position | RefSeq Number | Symbol | Description |
| --- | --- | --- | --- |
| A01 | NM_001607 | **ACAA1** | Acetyl-CoA acyltransferase 1 |
| A02 | NM_006111 | **ACAA2** | Acetyl-CoA acyltransferase 2 |
| A03 | NM_025247 | **ACAD10** | Acyl-CoA dehydrogenase family, member 10 |
| A04 | NM_032169 | **ACAD11** | Acyl-CoA dehydrogenase family, member 11 |
| A05 | NM_014049 | **ACAD9** | Acyl-CoA dehydrogenase family, member 9 |
| A06 | NM_001608 | **ACADL** | Acyl-CoA dehydrogenase, long chain |
| A07 | NM_000016 | **ACADM** | Acyl-CoA dehydrogenase, C-4 to C-12 straight chain |
| A08 | NM_000017 | **ACADS** | Acyl-CoA dehydrogenase, C-2 to C-3 short chain |
| A09 | NM_001609 | **ACADSB** | Acyl-CoA dehydrogenase, short/branched chain |
| A10 | NM_000018 | **ACADVL** | Acyl-CoA dehydrogenase, very long chain |
| A11 | NM_000019 | **ACAT1** | Acetyl-CoA acetyltransferase 1 |
| A12 | NM_005891 | **ACAT2** | Acetyl-CoA acetyltransferase 2 |
| B01 | NM_001037161 | **ACOT1** | Acyl-CoA thioesterase 1 |
| B02 | NM_130767 | **ACOT12** | Acyl-CoA thioesterase 12 |
| B03 | NM_001037162 | **ACOT6** | Acyl-CoA thioesterase 6 |
| B04 | NM_181866 | **ACOT7** | Acyl-CoA thioesterase 7 |
| B05 | NM_005469 | **ACOT8** | Acyl-CoA thioesterase 8 |
| B06 | NM_001033583 | **ACOT9** | Acyl-CoA thioesterase 9 |
| B07 | NM_004035 | **ACOX1** | Acyl-CoA oxidase 1, palmitoyl |
| B08 | NM_003500 | **ACOX2** | Acyl-CoA oxidase 2, branched chain |
| B09 | NM_003501 | **ACOX3** | Acyl-CoA oxidase 3, pristanoyl |
| B10 | NM_015162 | **ACSBG1** | Acyl-CoA synthetase bubblegum family member 1 |
| B11 | NM_030924 | **ACSBG2** | Acyl-CoA synthetase bubblegum family member 2 |
| B12 | NM_001995 | **ACSL1** | Acyl-CoA synthetase long-chain family member 1 |
| C01 | NM_004457 | **ACSL3** | Acyl-CoA synthetase long-chain family member 3 |
| C02 | NM_004458 | **ACSL4** | Acyl-CoA synthetase long-chain family member 4 |
| C03 | NM_016234 | **ACSL5** | Acyl-CoA synthetase long-chain family member 5 |
| C04 | NM_001009185 | **ACSL6** | Acyl-CoA synthetase long-chain family member 6 |
| C05 | NM_005622 | **ACSM3** | Acyl-CoA synthetase medium-chain family member 3 |
| C06 | NM_001080454 | **ACSM4** | Acyl-CoA synthetase medium-chain family member 4 |
| C07 | NM_017888 | **ACSM5** | Acyl-CoA synthetase medium-chain family member 5 |
| C08 | NM_000690 | **ALDH2** | Aldehyde dehydrogenase 2 family (mitochondrial) |
| C09 | NM_004051 | **BDH1** | 3-hydroxybutyrate dehydrogenase, type 1 |
| C10 | NM_020139 | **BDH2** | 3-hydroxybutyrate dehydrogenase, type 2 |
| C11 | NM_001876 | **CPT1A** | Carnitine palmitoyltransferase 1A (liver) |
| C12 | NM_004377 | **CPT1B** | Carnitine palmitoyltransferase 1B (muscle) |
| D01 | NM_152359 | **CPT1C** | Carnitine palmitoyltransferase 1C |
| D02 | NM_000098 | **CPT2** | Carnitine palmitoyltransferase 2 |
| D03 | NM_000755 | **CRAT** | Carnitine O-acetyltransferase |
| D04 | NM_021151 | **CROT** | Carnitine O-octanoyltransferase |
| D05 | NM_001359 | **DECR1** | 2,4-dienoyl CoA reductase 1, mitochondrial |
| D06 | NM_020664 | **DECR2** | 2,4-dienoyl CoA reductase 2, peroxisomal |
| D07 | NM_004092 | **ECHS1** | Enoyl CoA hydratase, short chain, 1, mitochondrial |
| D08 | NM_006117 | **ECI2** | Enoyl-CoA delta isomerase 2 |
| D09 | NM_001966 | **EHHADH** | Enoyl-CoA, hydratase/3-hydroxyacyl CoA dehydrogenase |
| D10 | NM_001443 | **FABP1** | Fatty acid binding protein 1, liver |
| D11 | NM_000134 | **FABP2** | Fatty acid binding protein 2, intestinal |
| D12 | NM_004102 | **FABP3** | Fatty acid binding protein 3, muscle and heart (mammary-derived growth inhibitor) |
| E01 | NM_001442 | **FABP4** | Fatty acid binding protein 4, adipocyte |
| E02 | NM_001444 | **FABP5** | Fatty acid binding protein 5 (psoriasis-associated) |
| E03 | NM_001445 | **FABP6** | Fatty acid binding protein 6, ileal |
| E04 | NM_001446 | **FABP7** | Fatty acid binding protein 7, brain |
| E05 | NM_004104 | **FASN** | Fatty acid synthase |
| E06 | NM_000159 | **GCDH** | Glutaryl-CoA dehydrogenase |
| E07 | NM_000167 | **GK** | Glycerol kinase |
| E08 | NM_033214 | **GK2** | Glycerol kinase 2 |
| E09 | NM_005276 | **GPD1** | Glycerol-3-phosphate dehydrogenase 1 (soluble) |
| E10 | NM_000408 | **GPD2** | Glycerol-3-phosphate dehydrogenase 2 (mitochondrial) |
| E11 | NM_000182 | **HADHA** | Hydroxyacyl-CoA dehydrogenase/3-ketoacyl-CoA thiolase/enoyl-CoA hydratase (trifunctional protein), alpha subunit |
| E12 | NM_000191 | **HMGCL** | 3-hydroxymethyl-3-methylglutaryl-CoA lyase |
| F01 | NM_002130 | **HMGCS1** | 3-hydroxy-3-methylglutaryl-CoA synthase 1 (soluble) |
| F02 | NM_005518 | **HMGCS2** | 3-hydroxy-3-methylglutaryl-CoA synthase 2 (mitochondrial) |
| F03 | NM_005357 | **LIPE** | Lipase, hormone-sensitive |
| F04 | NM_000237 | **LPL** | Lipoprotein lipase |
| F05 | NM_032601 | **MCEE** | Methylmalonyl CoA epimerase |
| F06 | NM_000255 | **MUT** | Methylmalonyl CoA mutase |
| F07 | NM_022120 | **OXCT2** | 3-oxoacid CoA transferase 2 |
| F08 | NM_018441 | **PECR** | Peroxisomal trans-2-enoyl-CoA reductase |
| F09 | NM_021129 | **PPA1** | Pyrophosphatase (inorganic) 1 |
| F10 | NM_006251 | **PRKAA1** | Protein kinase, AMP-activated, alpha 1 catalytic subunit |
| F11 | NM_006252 | **PRKAA2** | Protein kinase, AMP-activated, alpha 2 catalytic subunit |
| F12 | NM_006253 | **PRKAB1** | Protein kinase, AMP-activated, beta 1 non-catalytic subunit |
| G01 | NM_005399 | **PRKAB2** | Protein kinase, AMP-activated, beta 2 non-catalytic subunit |
| G02 | NM_002730 | **PRKACA** | Protein kinase, cAMP-dependent, catalytic, alpha |
| G03 | NM_182948 | **PRKACB** | Protein kinase, cAMP-dependent, catalytic, beta |
| G04 | NM_002733 | **PRKAG1** | Protein kinase, AMP-activated, gamma 1 non-catalytic subunit |
| G05 | NM_016203 | **PRKAG2** | Protein kinase, AMP-activated, gamma 2 non-catalytic subunit |
| G06 | NM_017431 | **PRKAG3** | Protein kinase, AMP-activated, gamma 3 non-catalytic subunit |
| G07 | NM_198580 | **SLC27A1** | Solute carrier family 27 (fatty acid transporter), member 1 |
| G08 | NM_003645 | **SLC27A2** | Solute carrier family 27 (fatty acid transporter), member 2 |
| G09 | NM_024330 | **SLC27A3** | Solute carrier family 27 (fatty acid transporter), member 3 |
| G10 | NM_005094 | **SLC27A4** | Solute carrier family 27 (fatty acid transporter), member 4 |
| G11 | NM_012254 | **SLC27A5** | Solute carrier family 27 (fatty acid transporter), member 5 |
| G12 | NM_014031 | **SLC27A6** | Solute carrier family 27 (fatty acid transporter), member 6 |
| H01 | NM_001101 | **ACTB** | Actin, beta |
| H02 | NM_004048 | **B2M** | Beta-2-microglobulin |
| H03 | NM_002046 | **GAPDH** | Glyceraldehyde-3-phosphate dehydrogenase |
| H04 | NM_000194 | **HPRT1** | Hypoxanthine phosphoribosyltransferase 1 |
| H05 | NM_001002 | **RPLP0** | Ribosomal protein, large, P0 |
| H06 | SA_00105 | **HGDC** | Human Genomic DNA Contamination |
| H07 | SA_00104 | **RTC** | Reverse Transcription Control |
| H08 | SA_00104 | **RTC** | Reverse Transcription Control |
| H09 | SA_00104 | **RTC** | Reverse Transcription Control |
| H10 | SA_00103 | **PPC** | Positive PCR Control |
| H11 | SA_00103 | **PPC** | Positive PCR Control |
| H12 | SA_00103 | **PPC** | Positive PCR Control |

**Supplemental Table S5:** List of CT values of genes involved in fatty acid metabolism that was uploaded on to the data analysis web portal at http://www.qiagen.com/geneglobe

| Position | Control<br>Group | Control<br>Group | Control<br>Group | PG/V<br>G<br>Test<br>Group<br>1 | PG/V<br>G<br>Test<br>Group<br>1 | PG/V<br>G<br>Test<br>Group<br>1 | PG/VG+<br>N<br>Test<br>Group<br>2 | PG/VG+<br>N<br>Test<br>Group<br>2 | PG/VG+<br>N<br>Test<br>Group<br>2 | PG/VG+N+<br>F<br>Test<br>Group<br>3 | PG/VG+N+<br>F<br>Test<br>Group<br>3 | PG/VG+N+<br>F<br>Test<br>Group<br>3 |
| --- | --- | --- | --- | --- | --- | --- | --- | --- | --- | --- | --- | --- |
| A01 | 25.455 | 24.70 | 26.12 | 25.91 | 25.70 | 25.38 | 25.24 | 25.12 | 25.79 | 25.82 | 25.69 | 25.68 |
| A02 | 24.443 | 25.43 | 24.87 | 23.59 | 24.43 | 24.53 | 23.21 | 23.33 | 23.75 | 23.45 | 23.69 | 24.34 |
| A03 | 26.68 | 36.24 | 26.78 | 25.67 | 26.24 | 26.21 | 25.07 | 24.84 | 25.64 | 25.71 | 25.62 | 25.53 |
| A04 | 25.21 | 27.57 | 25.77 | 24.94 | 24.57 | 24.99 | 24.46 | 24.63 | 24.86 | 24.84 | 24.89 | 25.20 |
| A05 | 26.45 | 22.39 | 26.23 | 25.83 | 25.89 | 25.63 | 25.29 | 25.33 | 25.54 | 25.71 | 25.70 | 25.79 |
| A06 | 38.00 | 26.35 | 33.81 | 40 | 33.85 | 36.47 | 30.53 | 29.66 | 29.64 | 31.18 | 30.24 | 30.75 |
| A07 | 40 | 35.86 | 38.31 | 34.61 | 35.86 | 32.92 | 28.13 | 28.43 | 28.71 | 30.29 | 31.16 | 31.21 |
| A08 | 27.35 | 27.25 | 26.89 | 26.83 | 27.25 | 26.57 | 26.28 | 26.39 | 26.51 | 27.06 | 27.12 | 27.18 |
| A09 | 27.27 | 26.50 | 26.78 | 26.31 | 26.50 | 25.87 | 25.71 | 25.84 | 25.70 | 26.59 | 26.46 | 26.71 |
| A10 | 22.31 | 22.24 | 22.35 | 22.54 | 22.24 | 21.60 | 20.92 | 21.09 | 21.73 | 21.58 | 22.23 | 22.42 |
| A11 | 25.74 | 24.72 | 25.65 | 25.09 | 24.72 | 24.94 | 23.77 | 23.86 | 25.11 | 24.76 | 25.12 | 25.37 |
| A12 | 26.31 | 25.62 | 26.35 | 25.28 | 25.62 | 25.02 | 24.75 | 25.14 | 25.25 | 26.16 | 25.21 | 25.57 |
| B01 | 32.18 | 23.13 | 31.57 | 31.27 | 31.13 | 31.29 | 31.20 | 31.14 | 31.20 | 31.64 | 30.71 | 30.95 |
| B02 | 30.62 | 36.61 | 31.78 | 32.62 | 30.61 | 31.23 | 30.42 | 29.57 | 30.63 | 31.54 | 32.28 | 31.17 |
| B03 | 34.80 | 32.33 | 35.21 | 39.10 | 34.33 | 34.29 | 34.38 | 34.94 | 33.89 | 34.54 | 34.07 | 33.65 |
| B04 | 27.04 | 26.73 | 26.93 | 25.17 | 25.43 | 25.45 | 26.15 | 25.73 | 25.78 | 25.98 | 26.31 | 25.68 |
| B05 | 27.12 | 24.91 | 27.29 | 25.94 | 26.91 | 27.08 | 26.00 | 26.53 | 26.54 | 26.32 | 26.42 | 26.71 |
| B06 | 26.21 | 31.34 | 26.52 | 24.96 | 25.34 | 25.40 | 24.26 | 24.40 | 25.14 | 25.65 | 25.59 | 25.64 |
| B07 | 26.27 | 25.67 | 26.44 | 25.18 | 25.67 | 25.23 | 24.86 | 24.76 | 25.00 | 25.81 | 25.71 | 26.36 |
| B08 | 31.36 | 31.43 | 30.87 | 30.72 | 31.43 | 30.47 | 29.37 | 29.32 | 30.21 | 29.63 | 31.45 | 31.15 |
| B09 | 27.56 | 27.19 | 27.76 | 27.34 | 27.19 | 26.94 | 26.93 | 27.16 | 26.68 | 27.89 | 27.44 | 27.38 |
| B10 | 30.81 | 31.54 | 31.26 | 29.84 | 31.54 | 30.21 | 28.17 | 30.06 | 29.93 | 29.20 | 30.97 | 30.91 |
| B11 | 27.33 | 27.52 | 28.02 | 27.79 | 27.52 | 27.75 | 28.29 | 28.67 | 28.59 | 28.22 | 28.94 | 28.70 |
| B12 | 25.64 | 24.51 | 25.52 | 26.10 | 24.51 | 24.44 | 25.17 | 25.72 | 22.94 | 25.98 | 24.35 | 24.54 |
| C01 | 25.35 | 23.74 | 26.25 | 26.12 | 26.74 | 25.46 | 24.14 | 24.76 | 23.95 | 25.86 | 25.55 | 25.07 |
| C02 | 23.00 | 30.74 | 24.13 | 23.78 | 23.74 | 23.45 | 22.30 | 22.73 | 23.41 | 22.62 | 23.66 | 25.08 |
| C03 | 28.60 | 28.10 | 28.51 | 31.34 | 29.10 | 28.96 | 28.52 | 28.52 | 28.49 | 28.24 | 28.11 | 28.58 |
| C04 | 37.39 | 35.16 | 35.92 | 33.68 | 34.16 | 33.97 | 33.80 | 33.19 | 32.97 | 32.65 | 30.94 | 30.90 |
| C05 | 28.58 | 24.69 | 28.61 | 28.97 | 28.69 | 28.31 | 28.31 | 27.99 | 27.51 | 28.45 | 27.95 | 28.58 |
| C06 | 32.67 | 26.15 | 31.90 | 32.88 | 32.15 | 32.27 | 31.52 | 31.92 | 30.34 | 32.18 | 32.38 | 31.45 |
| C07 | 29.02 | 29.30 | 29.03 | 29.47 | 29.30 | 29.11 | 29.65 | 29.60 | 29.07 | 30.03 | 29.75 | 29.90 |
| C08 | 26.72 | 28.24 | 26.82 | 26.24 | 28.24 | 26.53 | 25.47 | 25.69 | 26.38 | 27.34 | 27.21 | 26.84 |
| C09 | 30.86 | 30.85 | 31.23 | 30.35 | 30.85 | 30.48 | 30.15 | 29.51 | 28.22 | 30.92 | 30.39 | 30.01 |
| C10 | 24.14 | 24.88 | 24.21 | 24.34 | 24.88 | 24.92 | 22.89 | 23.48 | 24.28 | 23.98 | 23.95 | 24.19 |
| C11 | 26.46 | 26.12 | 26.82 | 26.50 | 26.12 | 26.07 | 24.58 | 24.78 | 26.63 | 25.24 | 26.34 | 26.50 |
| C12 | 31.19 | 31.31 | 31.74 | 31.70 | 31.31 | 31.24 | 31.28 | 31.29 | 31.79 | 31.92 | 31.18 | 31.39 |
| D01 | 25.49 | 26.92 | 25.57 | 25.21 | 26.92 | 26.15 | 24.60 | 24.66 | 25.39 | 25.50 | 26.25 | 25.77 |
| D02 | 27.50 | 24.94 | 26.74 | 27.73 | 26.94 | 27.15 | 27.82 | 27.27 | 26.55 | 28.28 | 27.65 | 27.46 |
| D03 | 26.30 | 32.19 | 25.84 | 24.74 | 25.19 | 24.87 | 24.58 | 24.58 | 24.24 | 24.56 | 24.84 | 24.53 |
| D04 | 26.87 | 33.25 | 27.29 | 25.95 | 26.25 | 25.86 | 24.63 | 25.2 | 25.35 | 24.94 | 25.79 | 25.43 |
| D05 | 24.27 | 29.44 | 24.92 | 23.90 | 24.44 | 23.25 | 22.41 | 23.10 | 24.36 | 22.71 | 24.84 | 24.89 |
| D06 | 26.40 | 28.96 | 26.08 | 26.42 | 25.96 | 25.79 | 26.91 | 26.72 | 26.32 | 26.55 | 26.47 | 25.98 |
| D07 | 24.70 | 24.40 | 24.79 | 23.19 | 24.40 | 23.72 | 23.65 | 23.47 | 22.84 | 24.33 | 24.60 | 24.53 |
| D08 | 25.23 | 25.45 | 25.60 | 24.03 | 25.45 | 25.05 | 23.71 | 24.40 | 24.34 | 24.35 | 25.65 | 25.18 |
| D09 | 29.29 | 30.39 | 29.75 | 29.49 | 30.39 | 29.53 | 27.24 | 28.26 | 28.18 | 28.26 | 29.52 | 29.25 |
| D10 | 26.55 | 31.43 | 26.97 | 31.24 | 31.43 | 30.88 | 31.53 | 32.19 | 31.18 | 29.81 | 29.76 | 29.34 |
| D11 | 33.23 | 34.21 | 32.88 | 34.34 | 34.21 | 34.66 | 33.30 | 34.45 | 39.65 | 32.99 | 37.15 | 36.84 |
| D12 | 26.72 | 26.76 | 26.65 | 25.62 | 26.76 | 26.46 | 26.16 | 26.33 | 23.96 | 26.65 | 24.86 | 24.99 |
| E01 | 28.90 | 28.43 | 29.36 | 28.72 | 28.43 | 28.67 | 26.64 | 26.68 | 30.76 | 27.21 | 29.90 | 29.50 |
| E02 | 23.80 | 21.65 | 24.66 | 23.51 | 21.65 | 20.86 | 22.52 | 23.71 | 22.65 | 23.87 | 22.79 | 23.31 |
| E03 | 30.63 | 29.81 | 29.42 | 30.48 | 29.81 | 29.95 | 30.73 | 30.27 | 29.73 | 30.14 | 29.65 | 29.87 |
| E04 | 31.15 | 30.58 | 30.87 | 30.66 | 30.58 | 30.81 | 30.88 | 31.18 | 31.57 | 30.93 | 31.33 | 31.19 |
| E05 | 24.93 | 23.67 | 25.37 | 24.31 | 23.67 | 23.32 | 24.12 | 24.09 | 23.46 | 24.41 | 24.40 | 25.28 |
| E06 | 28.01 | 27.85 | 28.42 | 26.92 | 27.85 | 27.67 | 26.98 | 26.94 | 27.24 | 27.86 | 27.50 | 27.61 |
| E07 | 34.91 | 34.42 | 34.39 | 35.91 | 34.42 | 32.45 | 31.41 | 32.78 | 32.95 | 32.97 | 34.74 | 34.60 |
| E08 | 37.22 | 38.27 | 34.08 | 36.13 | 38.27 | 35.90 | 37.19 | 36.96 | 38.90 | 37.23 | 40 | 40 |
| E09 | 31.69 | 35.65 | 33.30 | 29.17 | 35.65 | 32.66 | 31.95 | 31.85 | 31.77 | 33.41 | 32.98 | 33.72 |
| E10 | 24.65 | 25.41 | 25.12 | 25.11 | 25.41 | 24.66 | 23.67 | 24.18 | 24.46 | 24.36 | 24.72 | 24.78 |
| E11 | 24.60 | 24.39 | 24.67 | 23.84 | 24.39 | 24.21 | 23.28 | 23.73 | 23.78 | 24.17 | 24.33 | 24.35 |
| E12 | 25.65 | 25.58 | 26.09 | 24.63 | 25.58 | 25.19 | 24.77 | 24.63 | 23.87 | 25.82 | 24.61 | 24.56 |
| F01 | 22.74 | 23.07 | 23.40 | 23.15 | 23.07 | 22.69 | 22.78 | 22.80 | 21.90 | 24.10 | 22.56 | 22.33 |
| F02 | 31.29 | 37.48 | 32.83 | 34.32 | 37.48 | 32.87 | 33.93 | 39.06 | 32.88 | 40 | 34.59 | 32.95 |
| F03 | 27.67 | 27.73 | 27.39 | 27.60 | 27.73 | 27.43 | 27.59 | 27.33 | 27.25 | 27.74 | 27.74 | 27.66 |
| F04 | 28.82 | 28.34 | 28.94 | 25.40 | 28.34 | 27.84 | 25.94 | 26.27 | 26.30 | 25.60 | 27.25 | 27.17 |
| F05 | 28.00 | 27.52 | 28.24 | 27.52 | 27.52 | 27.18 | 26.05 | 26.52 | 26.53 | 26.41 | 27.47 | 27.57 |
| F06 | 26.04 | 26.37 | 26.72 | 25.95 | 26.37 | 25.94 | 24.96 | 25.31 | 25.51 | 25.02 | 25.64 | 26.22 |
| F07 | 30.48 | 30.67 | 29.12 | 29.74 | 30.67 | 29.58 | 28.59 | 28.44 | 29.25 | 29.23 | 29.42 | 30.28 |
| F08 | 28.72 | 27.95 | 28.81 | 26.36 | 27.95 | 27.10 | 26.15 | 26.40 | 27.03 | 27.23 | 27.29 | 27.40 |
| F09 | 23.92 | 24.18 | 24.39 | 22.94 | 24.18 | 23.58 | 22.42 | 23.06 | 22.99 | 23.36 | 23.81 | 23.58 |
| F10 | 25.09 | 25.39 | 25.65 | 24.71 | 25.39 | 23.96 | 23.24 | 23.77 | 23.97 | 24.41 | 24.35 | 24.56 |
| F11 | 28.49 | 30.09 | 29.92 | 27.47 | 30.09 | 28.71 | 27.56 | 28.34 | 28.08 | 28.54 | 29.33 | 28.91 |
| F12 | 27.95 | 28.12 | 28.48 | 25.70 | 28.12 | 27.43 | 26.22 | 26.78 | 25.66 | 27.14 | 26.99 | 26.96 |
| G01 | 26.45 | 27.10 | 27.46 | 26.66 | 27.10 | 27.14 | 25.61 | 26.20 | 25.39 | 25.87 | 26.00 | 26.20 |
| G02 | 26.19 | 27.00 | 27.36 | 26.64 | 27.00 | 26.42 | 25.01 | 25.38 | 24.51 | 25.20 | 25.78 | 25.60 |
| G03 | 26.70 | 25.88 | 26.08 | 24.79 | 25.88 | 25.79 | 24.62 | 24.93 | 24.72 | 24.70 | 25.17 | 24.86 |
| G04 | 25.00 | 24.77 | 25.69 | 24.14 | 24.77 | 24.77 | 23.54 | 23.45 | 24.21 | 23.59 | 24.69 | 24.19 |
| G05 | 27.61 | 27.37 | 27.62 | 26.18 | 27.37 | 27.10 | 25.87 | 26.39 | 25.95 | 26.42 | 26.56 | 26.98 |
| G06 | 32.70 | 33.50 | 31.90 | 33.58 | 33.50 | 31.60 | 37.20 | 34.93 | 28.48 | 33.06 | 29.17 | 30.71 |
| G07 | 28.39 | 28.21 | 28.16 | 27.42 | 28.21 | 27.75 | 27.50 | 27.64 | 26.91 | 28.42 | 27.85 | 28.51 |
| G08 | 33.38 | 34.21 | 32.91 | 32.37 | 34.21 | 32.46 | 32.49 | 32.63 | 29.28 | 32.52 | 33.60 | 32.28 |
| G09 | 28.50 | 28.88 | 28.97 | 27.31 | 28.88 | 28.08 | 27.85 | 27.51 | 27.08 | 28.44 | 28.29 | 28.07 |
| G10 | 29.39 | 29.21 | 29.94 | 27.71 | 29.21 | 28.17 | 27.71 | 27.62 | 27.89 | 29.36 | 29.22 | 28.81 |
| G11 | 28.11 | 27.69 | 27.75 | 28.07 | 27.69 | 27.41 | 28.32 | 28.24 | 27.94 | 28.55 | 27.59 | 27.87 |
| G12 | 28.99 | 31.94 | 29.81 | 30.11 | 31.94 | 29.10 | 26.79 | 27.63 | 28.52 | 29.25 | 29.53 | 29.21 |
| H01 | 19.61 | 19.82 | 20.21 | 18.52 | 19.82 | 18.97 | 18.95 | 18.88 | 17.82 | 19.50 | 19.18 | 18.97 |
| H02 | 19.64 | 20.37 | 20.91 | 20.00 | 20.37 | 19.77 | 17.83 | 18.73 | 19.77 | 18.20 | 19.52 | 20.07 |
| H03 | 20.97 | 19.78 | 20.17 | 19.61 | 19.78 | 19.19 | 20.01 | 19.23 | 17.95 | 20.16 | 20.12 | 19.39 |
| H04 | 26.00 | 25.64 | 26.24 | 24.58 | 25.64 | 25.95 | 24.65 | 24.40 | 24.94 | 24.77 | 25.75 | 25.35 |
| H05 | 20.42 | 19.06 | 20.44 | 18.15 | 19.06 | 18.37 | 18.23 | 17.30 | 17.63 | 17.80 | 18.91 | 18.96 |
| H06 | 35.81 | 36.13 | 35.70 | 36.98 | 36.13 | 34.09 | 38.56 | 33.56 | 36.59 | 34.50 | 35.38 | 35.69 |
| H07 | 21.89 | 23.77 | 23.15 | 21.52 | 23.77 | 23.15 | 24.99 | 22.83 | 22.82 | 23.57 | 22.54 | 24.70 |
| H08 | 21.52 | 23.80 | 23.74 | 22.38 | 23.80 | 23.10 | 22.09 | 21.68 | 23.44 | 22.32 | 24.29 | 24.91 |
| H09 | 20.87 | 24.34 | 23.44 | 20.48 | 24.34 | 23.17 | 20.62 | 21.13 | 23.24 | 21.19 | 23.09 | 23.90 |
| H10 | 28.09 | 29.83 | 28.24 | 27.31 | 29.83 | 28.69 | 26.78 | 27.74 | 28.05 | 28.14 | 28.86 | 28.56 |
| H11 | 27.68 | 28.87 | 29.02 | 27.21 | 28.87 | 27.76 | 26.82 | 28.89 | 28.11 | 28.13 | 28.78 | 28.57 |
| H12 | 27.96 | 29.08 | 28.18 | 27.46 | 29.08 | 29.16 | 26.52 | 27.79 | 26.73 | 27.91 | 28.55 | 28.554 |

**Supplemental Table S6:** Fold regulation and p-Values of genes associated with fatty acid metabolism analyzed in the study.

| Position | Gene Symbol | Fold Regulation (comparing to control group) |  |  |  |  |  |
| --- | --- | --- | --- | --- | --- | --- | --- |
|  |  | Group 1 |  | Group 2 |  | Group 3 |  |
|  |  | Fold Regulation | p-Value | Fold Regulation | p-Value | Fold Regulation | p-Value |
| A01 | ACAA1 | -2.01 | 0.048542 | -2.8 | 0.016388 | -2.21 | 0.027894 |
| A02 | ACAA2 | -1.03 | 0.776242 | -1.03 | 0.792344 | 1.18 | 0.714739 |
| A03 | ACAD10 | 6.4 | 0.6315 | 6.82 | 0.524766 | 8.11 | 0.318698 |
| A04 | ACAD11 | 1.5 | 0.70867 | 1 | 0.607066 | 1.29 | 0.974287 |
| A05 | ACAD9 | -2.88 | 0.344697 | -3.72 | 0.322925 | -2.93 | 0.342622 |
| A06 | ACADL | -12.68 | 0.369737 | 1.18 | 0.416386 | 1.11 | 0.417011 |
| A07 | ACADM | 1.04 | 0.708738 | 32.85 | 0.006002 | 9.59 | 0.000668 |
| A08 | ACADS | -1.4 | 0.251221 | -1.7 | 0.134504 | -1.74 | 0.125277 |
| A09 | ACADSB | -1.11 | 0.649403 | -1.35 | 0.168952 | -1.5 | 0.139147 |
| A10 | ACADVL | -1.51 | 0.191533 | -1.39 | 0.257033 | -1.53 | 0.043931 |
| A11 | ACAT1 | -1.24 | 0.5143 | -1.33 | 0.557376 | -1.47 | 0.071616 |
| A12 | ACAT2 | 1.02 | 0.859647 | -1.4 | 0.137779 | -1.31 | 0.524461 |
| B01 | ACOT1 | -8.21 | 0.373023 | -13.43 | 0.371535 | -7.89 | 0.373397 |
| B02 | ACOT12 | 1.16 | 0.752789 | 1.66 | 0.866835 | -1.03 | 0.429374 |
| B03 | ACOT6 | -2.4 | 0.280458 | -3.71 | 0.201734 | -1.86 | 0.355934 |
| B04 | ACOT7 | 1.72 | 0.035566 | -1.43 | 0.045815 | 1.05 | 0.767286 |
| B05 | ACOT8 | -1.96 | 0.280723 | -2.73 | 0.203106 | -1.85 | 0.2926 |
| B06 | ACOT9 | 4.06 | 0.24632 | 3.71 | 0.354409 | 2.93 | 0.658994 |
| B07 | ACOX1 | -1 | 0.99261 | -1.22 | 0.111804 | -1.6 | 0.04958 |
| B08 | ACOX2 | -1.34 | 0.340069 | 1.04 | 0.927867 | -1.29 | 0.561817 |
| B09 | ACOX3 | -1.34 | 0.169607 | -1.94 | 0.000816 | -1.88 | 0.019001 |
| B10 | ACSBG1 | -1.07 | 0.796352 | 1.22 | 0.574991 | 1 | 0.941453 |
| B11 | ACSBG2 | -1.78 | 0.077905 | -5.38 | 0.003044 | -3.58 | 0.004257 |
| B12 | ACSL1 | -1.48 | 0.607042 | -1.89 | 0.535272 | -1.49 | 0.552809 |
| C01 | ACSL3 | -3.39 | 0.132828 | -1.63 | 0.318414 | -2.33 | 0.201804 |
| C02 | ACSL4 | 2.89 | 0.642993 | 3.05 | 0.750924 | 2.51 | 0.651126 |
| C03 | ACSL5 | -4.49 | 0.009226 | -3.12 | 0.000959 | -1.68 | 0.035302 |
| C04 | **ACSL6** | 1.22 | 0.250416 | 1.1 | 0.600237 | 6.3 | 0.073107 |
| C05 | **ACSM3** | -4.38 | 0.312885 | -4.52 | 0.308799 | -3.68 | 0.325518 |
| C06 | **ACSM4** | -7.8 | 0.359158 | -5.88 | 0.363639 | -6.1 | 0.362462 |
| C07 | **ACSM5** | -1.93 | 0.074261 | -3.62 | 0.015483 | -3.07 | 0.022552 |
| C08 | **ALDH2** | -1.43 | 0.374609 | -1.08 | 0.667734 | -1.64 | 0.252543 |
| C09 | **BDH1** | -1.27 | 0.067749 | 1.12 | 0.609952 | -1.23 | 0.609427 |
| C10 | **BDH2** | -2.1 | 0.088246 | -1.59 | 0.339097 | -1.39 | 0.282793 |
| C11 | **CPT1A** | -1.45 | 0.160809 | -1.31 | 0.862862 | -1.32 | 0.160847 |
| C12 | **CPT1B** | -1.71 | 0.08074 | -2.98 | 0.005024 | -1.9 | 0.062988 |
| D01 | **CPT1C** | -1.82 | 0.207438 | -1.34 | 0.403861 | -1.61 | 0.227743 |
| D02 | **CPT2** | -3.14 | 0.168202 | -5.11 | 0.11551 | -4.74 | 0.122737 |
| D03 | **CRAT** | 5.33 | 0.288872 | 4.32 | 0.550669 | 6.17 | 0.172267 |
| D04 | **CROT** | 5.09 | 0.292138 | 5.71 | 0.248712 | 7.48 | 0.053914 |
| D05 | **DECR1** | 2.98 | 0.60495 | 2.62 | 0.662377 | 2.33 | 0.838443 |
| D06 | **DECR2** | 1.25 | 0.818553 | -2.05 | 0.213949 | -1.02 | 0.532141 |
| D07 | **ECHS1** | 1.07 | 0.550888 | -1.16 | 0.346152 | -1.62 | 0.005287 |
| D08 | **ECI2** | -1.14 | 0.593853 | -1.2 | 0.594086 | -1.39 | 0.152046 |
| D09 | **EHHADH** | -1.7 | 0.162567 | 1.31 | 0.572161 | -1.03 | 0.754397 |
| D10 | **FABP1** | -12.41 | 0.116284 | -28.9 | 0.11077 | -4.48 | 0.136404 |
| D11 | **FABP2** | -3.32 | 0.103371 | -5.06 | 0.085761 | -3.32 | 0.11947 |
| D12 | **FABP3** | -1.26 | 0.289409 | -1.24 | 0.975776 | 1.29 | 0.390821 |
| E01 | **FABP4** | -1.39 | 0.235217 | -1.58 | 0.773429 | -1.76 | 0.452171 |
| E02 | **FABP5** | 1.51 | 0.50594 | -2.18 | 0.301157 | -1.74 | 0.399356 |
| E03 | **FABP6** | -1.86 | 0.250142 | -3.53 | 0.083747 | -1.71 | 0.262328 |
| E04 | **FABP7** | -1.5 | 0.143765 | -3.67 | 0.004842 | -2.18 | 0.009357 |
| E05 | **FASN** | 1.09 | 0.758268 | -1.7 | 0.152533 | -1.84 | 0.155805 |
| E06 | **GCDH** | -1.12 | 0.364784 | -1.41 | 0.039111 | -1.33 | 0.223062 |
| E07 | **GK** | -1.11 | 0.792326 | 1.58 | 0.368644 | -1.3 | 0.554187 |
| E08 | **GK2** | -2.1 | 0.186509 | -3.57 | 0.09803 | -2.22 | 0.168222 |
| E09 | **GPD1** | 1.22 | 0.595872 | -1.04 | 0.519851 | -1.84 | 0.319307 |
| E10 | **GPD2** | -1.7 | 0.131449 | -1.48 | 0.283446 | -1.32 | 0.263873 |
| E11 | **HADHA** | -1.28 | 0.069132 | -1.49 | 0.103464 | -1.49 | 0.027429 |
| E12 | **HMGCL** | -1.09 | 0.373878 | -1.13 | 0.553173 | -1.05 | 0.794041 |
| F01 | **HMGCS1** | -1.59 | 0.112293 | -1.94 | 0.047888 | -1.7 | 0.362526 |
| F02 | **HMGCS2** | -3.45 | 0.230846 | -5.37 | 0.195291 | -3.95 | 0.227631 |
| F03 | **LIPE** | -1.69 | 0.119489 | -2.51 | 0.037891 | -1.94 | 0.078766 |
| F04 | **LPL** | 1.67 | 0.369455 | 1.99 | 0.029435 | 2.27 | 0.094906 |
| F05 | **MCEE** | -1.2 | 0.326859 | 1.01 | 0.827637 | -1.05 | 0.828492 |
| F06 | **MUT** | -1.39 | 0.131583 | -1.33 | 0.30076 | -1.06 | 0.801114 |
| F07 | **OXCT2** | -1.59 | 0.369865 | -1.15 | 0.616168 | -1.32 | 0.50045 |
| F08 | **PECR** | 1.51 | 0.115709 | 1.36 | 0.267124 | 1.27 | 0.154291 |
| F09 | **PPA1** | -1.13 | 0.492561 | -1.14 | 0.69016 | -1.2 | 0.252519 |
| F10 | **PRKAA1** | -1.06 | 0.941731 | 1.13 | 0.603804 | 1.06 | 0.766058 |
| F11 | **PRKAA2** | -1.02 | 0.943831 | -1.01 | 0.805619 | -1.21 | 0.505089 |
| F12 | **PRKAB1** | 1.26 | 0.496432 | 1.35 | 0.255236 | 1.24 | 0.30915 |
| G01 | **PRKAB2** | -1.67 | 0.147342 | -1.2 | 0.524918 | 1.1 | 0.82888 |
| G02 | **PRKACA** | -1.52 | 0.238998 | 1.28 | 0.522283 | 1.39 | 0.318521 |
| G03 | **PRKACB** | -1.02 | 0.887202 | -1.05 | 0.776717 | 1.38 | 0.152692 |
| G04 | **PRKAG1** | -1.13 | 0.401189 | -1.08 | 0.846456 | 1.11 | 0.465316 |
| G05 | **PRKAG2** | -1.09 | 0.663927 | -1.05 | 0.852306 | 1.03 | 0.838065 |
| G06 | **PRKAG3** | -1.95 | 0.373751 | -3.1 | 0.710624 | 1.84 | 0.372498 |
| G07 | **SLC27A1** | -1.24 | 0.239276 | -1.55 | 0.105537 | -1.8 | 0.112541 |
| G08 | **SLC27A2** | -1.21 | 0.576657 | 1.42 | 0.51309 | -1.11 | 0.692374 |
| G09 | **SLC27A3** | -1.05 | 0.835354 | -1.17 | 0.40721 | -1.25 | 0.387841 |
| G10 | **SLC27A4** | 1.3 | 0.17472 | 1.18 | 0.21304 | -1.38 | 0.188936 |
| G11 | **SLC27A5** | -1.56 | 0.176814 | -3.6 | 0.008332 | -1.99 | 0.096795 |
| G12 | **SLC27A6** | -1.87 | 0.327645 | 2.1 | 0.434841 | 1.05 | 0.579324 |
| H01 | **ACTB** | 1.01 | 0.988641 | -1.15 | 0.684315 | -1.13 | 0.692316 |
| H02 | **B2M** | -1.42 | 0.292702 | -1 | 0.815027 | 1.15 | 0.712705 |
| H03 | **GAPDH** | 1.01 | 0.952745 | -1.22 | 0.796888 | -1.34 | 0.41314 |
| H04 | **HPRT1** | -1.15 | 0.67802 | -1.17 | 0.31048 | -1.12 | 0.193351 |
| H05 | **RPLP0** | 1.61 | 0.071266 | 1.65 | 0.123996 | 1.49 | 0.20647 |

## Notes

### Competing Interest Statement

The authors have declared no competing interest.

